# Role of Fibronectin in Postnatal Skeletal Development

**DOI:** 10.1101/2024.02.01.578475

**Authors:** Neha E. H. Dinesh, Nissan Baratang, Justine Rosseau, Ronit Mohapatra, Ling Li, Ramshaa Mahalingam, Kerstin Tiedemann, Philippe M. Campeau, Dieter P. Reinhardt

## Abstract

Fibronectin (FN) is a ubiquitous matrix glycoprotein essential for the physiological development of various tissues. Mutations in FN cause a form of skeletal dysplasia, emphasizing the importance of FN in cartilage and bone development. However, the relevance and functional role of FN during skeletal development remains elusive. We employed conditional knockout mouse models for the cellular FN isoform in cartilage (cFNKO), the plasma FN isoform in hepatocytes (pFNKO), and a double knockout (FNdKO) to determine the relevance of these two principal FN isoforms in postnatal skeletal development spanning from P1 to 2 months of age.

We identified a unique topological FN deposition pattern in the mouse limb with prominent levels at the resting and hypertrophic chondrocyte zones and in the trabecular bone. Circulating pFN did not enter the growth plate and was limited to the primary ossification center, whereas cartilage-specific cFN was detected as the major isoform in epiphyseal cartilage. Deletion of either one of the isoforms in single knockouts (cFNKO or pFNKO) only led to subtle changes in some of the analyzed parameters. Complete ablation of both cFN in the growth plate and circulating pFN in plasma resulted in significantly reduced postnatal body weight, body length, and bone length in the FNdKO mice. Assessment of the FNdKO adult bone microarchitecture using micro-CT revealed significantly reduced trabecular bone volume, trabecular network, bone mineral density, and increased bone marrow adiposity. Analysis of chondrogenesis in FNdKO mice showed changes in the proliferating and hypertrophic growth plate zones, consistent alterations in chondrogenic markers such as collagen type II and type X, reduced apoptosis of hypertrophic chondrocytes, and downregulation of bone formation markers. FNdKO mice also displayed decreased levels of transforming growth factor-β1 (TGFβ1) and downstream phospho-AKT levels, which are critical regulators of chondrogenesis and bone formation.

In conclusion, the data demonstrate that FN is essential for proper chondrogenesis and postnatal bone development. Simultaneous deletion of both FN isoforms in the developing cartilage leads to critical TGFβ-mediated alterations in chondrogenic differentiation, resulting in bone and skeletal defects.

**Significance/Highlights:**

- FN is highly expressed during mouse limb development with increased deposition in resting and hypertrophic chondrocyte zones and the primary ossification center.
- Cartilage-specific cFN and circulating pFN are distinctly distributed during embryonic and postnatal bone development, with chondrocyte-specific cFN present in the growth plate and pFN limited to trabecular bone and the bone marrow.
- Deletion of both cFN and pFN leads to reduced bone growth during early postnatal development.
- Deletion of cFN and pFN leads to reduced trabecular bone formation, bone mineralization, and increased bone marrow adiposity in 2-month adult mice.
- Absence of both FN isoforms in the FNdKO mouse model leads to altered chondrogenesis and reduced bone formation.
- FN regulates chondrogenesis via TGFβ-mediated phospho-AKT signaling.

## Introduction

Skeletal development in vertebrates begins during embryogenesis by condensation of mesenchymal stem cells (MSCs) originating from cranial neural crest cells, somites, and the lateral plate mesoderm ^1^. Neural crest and somite-derived MSCs form the axial skeleton by intramembranous ossification. The appendicular skeleton comprising the long bones develops exclusively from lateral plate mesoderm-derived MSCs by endochondral ossification ^2^. During this process, MSCs differentiate into chondrocytes that proliferate and mature into a hypertrophic state, forming the growth plate in the epiphyseal cartilage. The growth plate regulates the formation and growth of the appendicular skeleton, including long bones, during embryonic and postnatal development. Chondrocytes in growth plates are arranged in morphologically distinct zones with functional relevance to cartilage and skeletal development ^3^. The uppermost zone consists of resting chondrocytes, followed by a zone characterized by collagen type II-expressing columnar proliferating chondrocytes. These chondrocytes further differentiate into the pre-hypertrophic and mature hypertrophic chondrocytes that express collagen type X ^4^. These hypertrophic chondrocytes can then produce matrix-degrading metalloproteases (MMP-13,-9,-3) and undergo apoptosis ^5^ or can transdifferentiate into osteoblasts, osteocytes, and bone marrow adipocytes ^6,7^. Depending on the developmental time point, the hypertrophic cells contribute up to 40-70% of osteoblasts in the developing trabecular bone, thereby facilitating the formation of mineralized bone ^6,8–10^. Defects in extracellular matrix proteins alter this process of endochondral ossification in humans, leading to a wide range of skeletal pathologies ^11–18^. Spondylometaphyseal dysplasia associated with mutations in the matrix protein fibronectin (FN) represents a recently discovered skeletal pathology due to alterations in the growth plate ^19–22^.

FN is a multifunctional matrix glycoprotein present from the beginning of embryogenesis in all major vertebrate organ systems ^23,24^. FN is essential for survival with principal roles in embryonic neuronal patterning and cardiovascular development ^25,26^. FN is encoded by the fibronectin gene (*FN1*), located on chromosome 2 in humans and chromosome 1 in mice, and is transcribed into a ∼8kb mature mRNA ^27,28^. The pre-mRNA undergoes alternative splicing primarily of exons coding for three major domains, the extra domain A (EDA) and B (EDB), and the variable domain V to produce different FN isoforms. Overall, there are 27 reported human FN transcripts *(Ensemble GRCh38:CM000664.2)* and 17 transcripts *(Ensemble GRCm39:CM000994.3)* in the mouse. These FN isoforms are broadly classified as cellular FN (cFN), which form the insoluble fibers in the matrix and are produced by all matrix-producing cells **(Supp. Figure 1A)**. The other major isoform is the plasma FN (pFN), expressed only by hepatocytes in the liver **(Supp. Figure 1A)**. This isoform circulates in the blood and can enter several tissues, such as the heart, kidney, brain, and lung ^29,30^. The two isoforms sometimes have distinct and compensatory functions in various organ systems and pathological conditions ^31,32^. For example, we previously demonstrated that both cFN and pFN are required for postnatal development and maintenance of the aortic wall ^33^. Additionally, we identified in that study that in the absence of aortic smooth muscle-specific cFN, pFN can enter the aortic wall to compensate for cFN and safeguard the aortic structure and function.

In the context of the skeletal system, FN was shown to promote chondrocyte differentiation in several *in vitro* studies using chick limb buds and other cell culture models (*refer to review* ^34^). In adult mice, the pFN circulating isoform enters mature bone and is involved in bone matrix regulation ^35^. Notably, the identification of autosomal dominant mutations in FN as a cause for spondylometaphyseal dysplasia of the corner fracture type (SMDCF; OMIM #184255) suggested that FN is required for proper skeletal development in humans ^20–22^. Individuals with SMDCF display a wide range of skeletal anomalies, such as irregular ossification of the metaphysis that appears as corner fractures in X-ray radiographs, widened metaphyses, short stature caused by reduced bone growth, scoliosis, coxa vara, and fractures, among others. Additionally, our recent work demonstrated that SMDCF-causing mutations in FN impaired stem cell differentiation into chondrocytes ^19^. These data strongly suggest an essential role of FN during cartilage and bone growth. The skeletal system undergoes constant remodeling during embryogenesis and early postnatal (P) growth, gradually transitioning from cartilage to mature mineralized bone ^2^. It is not clear how FN is expressed and assembled during this process. Moreover, the importance of the cartilage-specific cFN and circulating pFN during postnatal skeletal development is not known. In this study, we investigate these aspects by employing conditional mouse models to knockout cartilage-specific cFN (cFNKO), hepatocyte-specific pFN (pFNKO), and both FN isoforms (FNdKO). We identified that deletion of both cFN and pFN in the FNdKO mouse model alters the skeletal growth in these mice from P1 to 2 months. Absence of both FN isoforms leads to impaired chondrogenesis in the growth plate and reduced formation of trabecular bone, which was associated with a reduction in transforming growth factor-β1 (TGFβ1) and phospho-AKT levels. Overall, the findings from this study elucidate the functional importance of the two FN isoforms in vertebrate postnatal skeletal development.

## Results

### FN has a distinct deposition pattern in mouse cartilage during gestation and postnatal cartilage development

To test the importance of FN during embryonic and postnatal skeletal development, we first analyzed the presence and deposition pattern of FN during various development phases in Fn1 floxed mice (Fn1*^flox/flox^*) ^32^, representing the control mice in this study. Limb development in mice begins around E9.0 when the mesenchyme buds are formed, and the organization of limb structures is visible by E12.5 ^36^. Hence, we performed indirect immunostaining for total FN (cFN and pFN) on whole embryo sections starting at embryonic day E12.5 to 2 months of age and did an overall qualitative assessment of the key features pertaining to FN deposition in cartilage and bone **(Figure 1A)**. FN is present early during embryogenesis, and as expected, we observed an intense staining for FN in various regions of the whole tissue section. High levels of FN were localized in and around the developing vertebral bodies **(Figure 1A; arrowheads)**. By E16.5, we observed that the limbs had developed a growth plate with distinct chondrocyte distribution zones and showed a developed primary ossification center **(Figure 1B; left panel; arrowheads)**. Immunostaining for total FN on tibiae at E16.5 showed increased FN deposition within the growth plate in the resting chondrocyte zone and the hypertrophic zone and reduced levels in the proliferating zone. Additionally, we noted high FN levels in the primary ossification center comprising the trabecular bone. Further, the cartilage in the developing digits also showed substantial FN deposition **(Figure 1B; right panel; arrowheads)**. At postnatal day 1 (P1), the tibiae showed more pronounced FN deposition in the resting zone and the center of the growth plate above the hypertrophic zone, which was characteristic for this time point **(Figure 1B; arrowhead)**. Relative to E16.5, the hypertrophic zone at P1 showed much lower levels of localized FN. By P15, we observed the development of the secondary ossification center in the resting zone of the growth plate with localization of FN around the secondary ossification center. At this point, FN was still present in the growth plate and the trabecular bone in the primary ossification center. With further development of the primary and secondary ossification center, the length of the growth plate reduces over time to become a narrow layer, which can be noted in P30 and 2-month-old tibiae **(Figure 1B)**. At P30, FN is primarily present in the hypertrophic and the trabecular zones. In a fully formed mature 2-month-old bone, the various zones of the growth plate are not clearly distinguishable. Still, we observed that most of the FN in bone is localized in the growth plate and the trabecular bone, and very little FN is present in the bone marrow **(Figure 1B)**. These findings demonstrate that FN is present early in the developing mouse limbs and that the deposition of FN in cartilage has a unique topological pattern.

**Figure 1.**
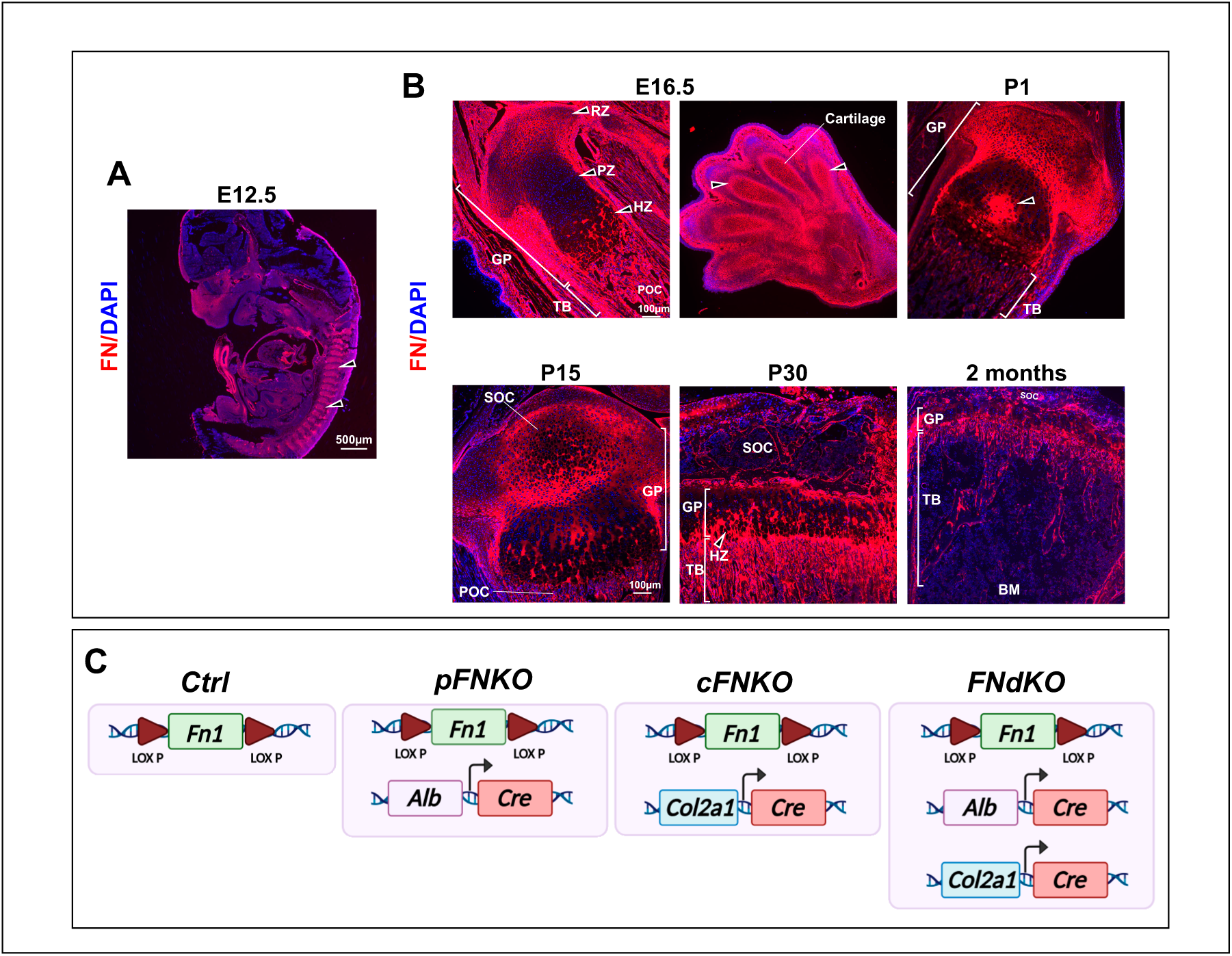
Deposition of FN in cartilage during embryonic and postnatal bone development. **(A)** Immunofluorescence staining of total FN (red), including cFN and pFN in a whole control mouse embryo section at E12.5. Note the intense staining for FN in the developing cartilage of vertebral bodies (arrowheads). One representative image for one mouse is shown of a total of n=11 mice analyzed. The scale bar represents 500 μm. **(B)** Immunofluorescence staining for total FN in mouse tibiae or fore paw from E16.5 to 2 months of age. For E16.5, the left image represents a tibia with total FN staining within the growth plate (GP) throughout the resting chondrocyte zone (RZ), proliferating chondrocyte zone (PZ), hypertrophic zone (HZ), trabecular bone (TB) and the primary ossification center (POC). The right image shows total FN staining in the fore paw in and around cartilage (arrowheads) (n=12). Images at P1, P15, P30, and 2-month-old mouse tibiae show total FN immunostaining in the GP, TB, secondary ossification center (SOC), POC, and bone marrow (BM). Representative images are shown from a total number of analyzed mice at P1 (n=18), P15 (n=7), P30 (n=9), and 2 months (n=9). The scale bar represents 100 μm. DAPI (blue) represents the cell nuclei in all images. **(C)** Schematic genotype overview of the transgenic mouse models employed in the study. Mice homozygous for the fibronectin gene (Fn1) flanked by lox P sites (Fn1(flox/flox), as described in ^32^) were used as controls (Ctrl) in all experiments. These mice crossed with a heterozygous transgene for Cre recombinase expression under the promoter of the albumin gene (Fn1(flox/flox); Alb(Cre/+) ^32^) represent plasma FN knockout mice (pFNKO). Homozygous floxed Fn1 mice crossed with a heterozygous transgene for Cre recombinase expression under the promoter of the collagen type II gene (Fn1(flox/flox); Col2a1(Cre/+)) represent cellular FN knockout mice (cFNKO). Mice expressing both Cre transgenes in homozygous floxed Fn1 mice represent FN double knockout mice (FNdKO). Note that the schematic is simplified, not showing both genotype alleles.

### Generation of conditional knockout mouse models for FN isoforms in cartilage

To further investigate the relevance of FN in postnatal cartilage and bone development, we employed the Cre-lox system to generate conditional knockout mice for the two principal isoforms of FN and completely ablate FN from the developing cartilage and bone **(Figure 1C)**. We crossed the *Fn1* gene floxed mice (Ctrl; Fn*^flox/flox^*) with transgenic mice expressing Cre recombinase under the regulation of the *Col2a1* promoter *(Col2a1-Cre/+)* to generate a cartilage-specific cFN knockout model (cFNKO; Fn*^flox/flox^*; *Col2a1-Cre/+)*. To generate the plasma FN knockout (pFNKO; Fn*^flox/flox^*; *Alb-Cre/+)*, Fn*^flox/flox^* mice were crossed with transgenic mice expressing Cre recombinase under the *Alb* promoter *(Alb-Cre/+)* as performed previously ^33^. The cFNKO and pFNKO were crossed to generate the FN double knockout model (FNdKO; Fn*^flox/flox^*; *Alb-Cre/+; Col2a1-Cre/+)* **(Figure 1C)**. *Col2a1* is expressed in mouse cartilage starting around E12.5, and the *Col2a1-Cre/+* model shows documented efficient Cre-mediated recombination in cartilage as early as E14.5 ^37^. The Alb-Cre transgene expression begins in the liver at E15.5 and is fully active by postnatal day 3 ^38^. Since albumin and pFN are exclusively expressed by hepatocytes in the liver, one can expect pFN to be deleted during both the fetal and neonatal stages. In this study, we focused on postnatal skeletal development, and hence, we analyzed the deletion of pFN at P1 and in adult mice at 2 months. As expected, pFN levels were dramatically reduced at P1 in the pFNKO **(Supp. Figure 1B)** and were completely absent at 2 months of age relative to the control mice **(Supp. Figure 1C)**.

### Plasma FN does not enter the growth plate during skeletal development, and cellular FN is the major isoform in cartilage

We next analyzed total FN levels in the cartilage of the control and FN knockout mice starting at E12.5. Imaging of E12.5 embryos showed no obvious morphological defects **(Figure 2A)**. As the limbs are not well developed at this age, we focused on cartilage forming the vertebral bodies. FN staining was present in and around the vertebral bodies in control and pFNKO mice **(Figure 2B; arrowheads).** However, we noted a reduction in total FN in the vertebral cartilage for FNdKO and cFNKO **(Figure 2B; arrowheads)**. From E16.5 to 2 months, we analyzed the growth plates in tibiae. As described above, in E16.5 to 2-month-old control mice tibia, FN was prominently present in the growth plate, with a distinct pattern of high levels of FN in the resting zone, hypertrophic zone, and trabecular bone **(Figure 2C-G)**. For the FNdKO mice at E16.5, FN was absent in the growth plate with minor staining near the trabecular bone region. However, from P1 to 2 months of age, FNdKO tissues were completely devoid of FN in the growth plate and trabecular bone **(Figure 2C-G)**. The single knockout mice, however, displayed a distinct FN staining pattern. Like FNdKO, the cFNKO was devoid of FN in the growth plate at all time points, demonstrating that cFN is the major isoform present in the cartilage during embryonic and postnatal cartilage development **(Figure 2C-G; arrowheads)**. However, FN was present in the developing trabecular bone and primary ossification center of the cFNKO tibiae **(Figure 2C-G)**, originating likely from the vasculature ingrowth that facilitates infiltration of pFN into the bone ^35^. An almost opposite FN distribution pattern was noted in pFNKO tibiae, where FN was always present within the analyzed time frame in the growth plate and remarkably reduced in the trabecular bone and primary ossification center **(Figure 2C-G; arrowheads).** These data demonstrate that both cFN and pFN have a unique physiological distribution in the developing bone, with cFN primarily present in the growth plate and pFN limited to trabecular bone, primary and secondary ossification centers **(Figure 2B-G; arrowheads)**.

**Figure 2.**
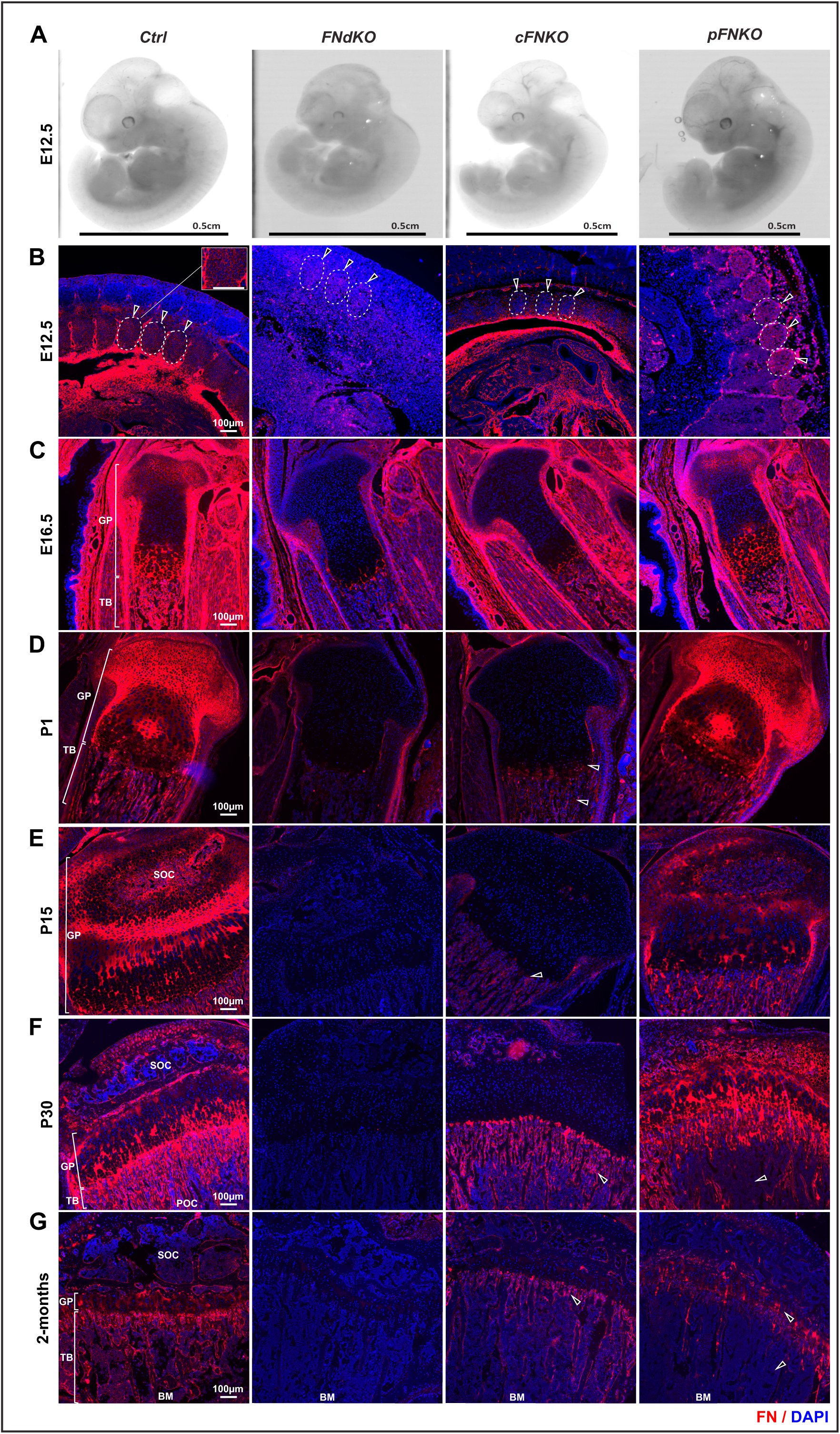
Analysis of FN in the transgenic FN mouse models during embryonic and postnatal cartilage development. **(A)** Gross images of E12.5 Ctrl and FN knockout embryos. The scale bar represents 0.5 cm **(B)** Immunostaining for total FN (red) at E12.5 on whole embryo sections. Note the FN deposition in developing vertebral bodies. The total number of mice analyzed were for ctrl (n=11), FNdKO (n=6), cFNKO (n=3), and pFNKO (n=15). **(C-G)** Total FN immunostaining on E16.5 to 2-month-old tibia sections. The total number of mice analyzed with similar stating patterns were for the Ctrl (n=7-18), FNdKO (n=6-15), cFNKO (n=3-19), and pFNKO (n=6-15). The scale bar represents 100 μm. DAPI (blue) was used as a nuclear counter stain. Growth plate, GP; Trabecular bone, TB; Primary ossification center, POC; Secondary ossification center, SOC; Bone marrow, BM.

### Deletion of cellular and plasma FN leads to reduced skeletal and long bone growth

FN is essential for mesenchymal condensation and chondrogenesis, which is well demonstrated in cell culture by us and others ^19,39^. Based on this existing knowledge, it was surprising to note that deletion of FN isoforms did not have detrimental effects on the survival of the FN knockout mice. To further investigate the consequence of FN isoform deletion on postnatal bone growth, we performed alcian blue and alizarin red whole-mount skeletal staining on P1 pup skeletons **(Figure 3)**. The FNdKO was smaller than the control and single knockout pups **(Figure 3A)**. No obvious defects in the skull **(Figure 3B)** or the ribcage **(Supp. Figure 2A)** were noted in any knockout mice using this technique. However, the length of bones comprising the forelimbs and hindlimbs was reduced for the FNdKO **(Figure 3C-D)**. Quantification of the total bone length **(Figure 3E)** and the mineralized bone length **(Supp. Figure 2B)** for the tibia, femur, humerus, radius, and ulna were significantly reduced in the FNdKO compared to control mice. Analysis of vertebral columns revealed the absence of ossification in the FNdKO vertebral bodies at the upper cervical vertebrae **(Supp. Figure 3A)** and reduced length of the lumber vertebrae from L1 to L6 **(Supp. Figure 3B)**.

**Figure 3.**
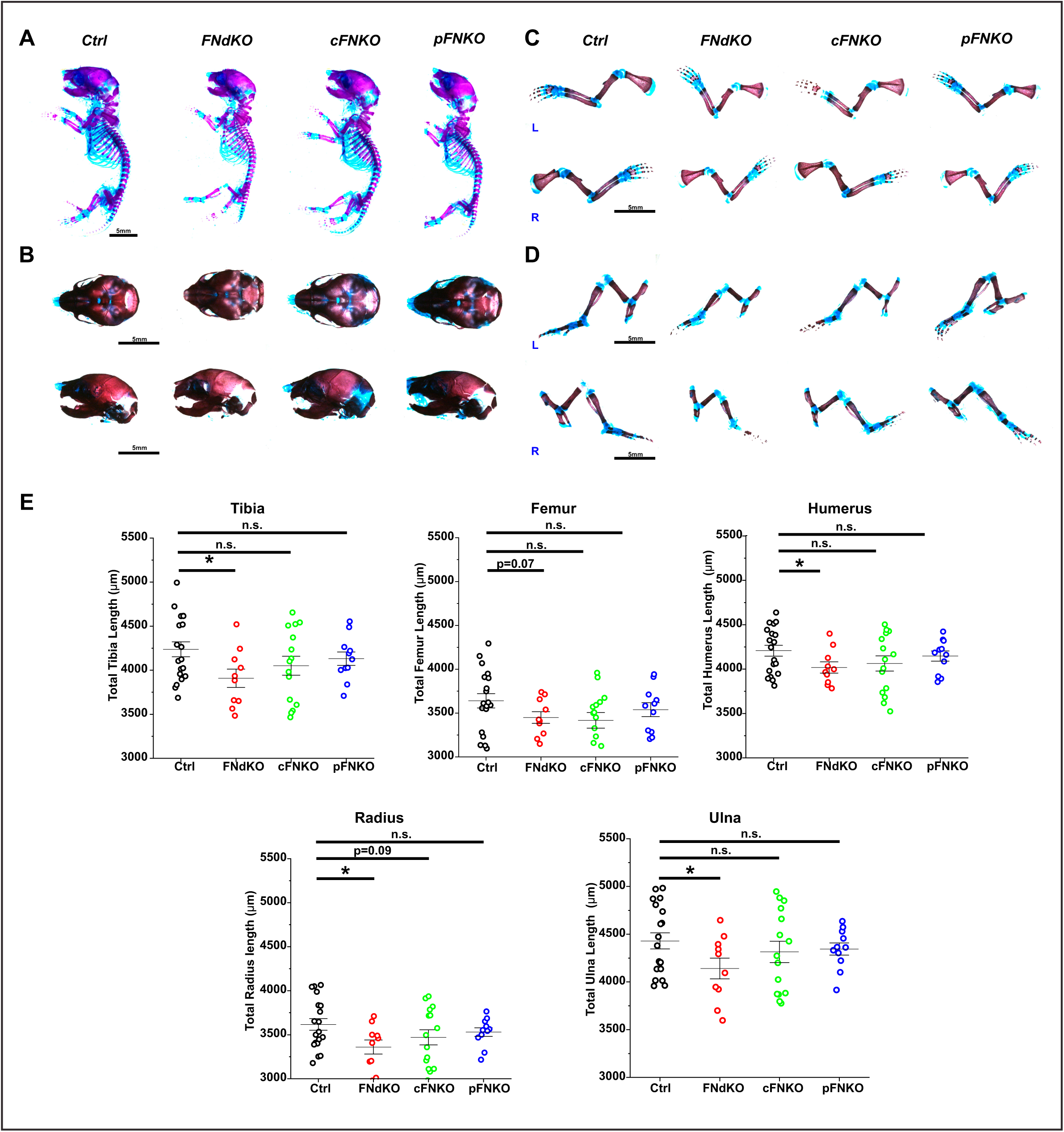
Whole-mount skeletal staining analysis at P1. **(A)** Gross images of P1 skeletons stained with Alcian blue (blue, cartilage) and alizarin red (red, mineralized bone). **(B)** Top and side views of skulls from control and FN knockout pups **(C-D)**. Arms and legs of P1 control and FN knockout pups (L represents left and R represents right). **(E)** Quantification of total bone length of the tibia, femur, humerus, radius, and ulna. Note the significantly reduced total bone length in the FNdKO. Each data point represents one pup. Error bars represent the standard error of the mean. Ctrl (n=19); FNdKO (n=10); cFNKO (n=15); pFNKO (n=11). The scale bar represents 5 mm. * Represents a p-value of <0.05. “n.s.” indicates a non-significant p-value.

FN knockout mice at P1 or P15 did have similar body weights **(Supp. Figure 4A,B)**. However, from P30 onwards, FNdKO mice displayed a significant reduction in total body length, body weight, and body weight normalized to total body length **(Supp. Figure 4C)**. These results suggest that the decrease in total body weight became more pronounced as postnatal skeletal development proceeded with age. Further, the FNdKO at P30 exhibited a significantly reduced length of the tibia, femur, humerus, and radius/ulna **(Supp. Figure 4D)**. The decreased bone length at P1 and P30 demonstrates that the alterations in long bone growth occur early and persist during postnatal development upon simultaneous deletion of cFN and pFN.

### Deletion of cellular and plasma FN leads to altered trabecular bone with increased bone marrow adiposity

To determine whether the deletion of FN isoforms in cartilage affects the formation of mature bone, we analyzed tibiae of 2-month-old male mice. The FNdKO exhibited significantly reduced overall body length and body weight compared to control mice, but the single knockout mice did not differ **(Figure 4A-B).** To analyze the trabecular bone, we performed micro-computed tomography (micro-CT) on tibia samples **(Figure 4C).** As observed at other time points, the length of the long bones, tibia, femur, radius/ulna, and humerus were all reduced in the FNdKO relative to controls **(Figure 4D)**. The tibia and radius in cFNKO mice were also significantly shorter, but not for the pFNKO samples. Additionally, the trabecular bone volume and bone volume to tissue volume ratio were significantly reduced for FNdKO but not for cFNKO and pFNKO **(Figure 4C,E)**. Other bone parameters such as bone surface density, bone mineral density, trabecular separation and number were consistently altered only in the double knockout mice **(Figure 4E-F)**. Bone porosity and trabecular thickness remained unaltered in all FN knockout mice **(Figure 4F)**. Bone marrow adiposity increases with age in both mice and humans, and increased adipocytes in the bone marrow indicate poor bone quality ^40^. Hence, to analyze this aspect in the FN knockout mice, we employed histological analysis of 2-month-old tibiae sections by H&E staining **(Supp. Figure 5A)**. The FNdKO exhibited increased bone marrow adipocytes, but not the single knockout mice. This suggests that pFN and cFN together play an essential and concerted role in regulating bone marrow adiposity **(Supp. Figure 5A,B)**.

**Figure 4.**
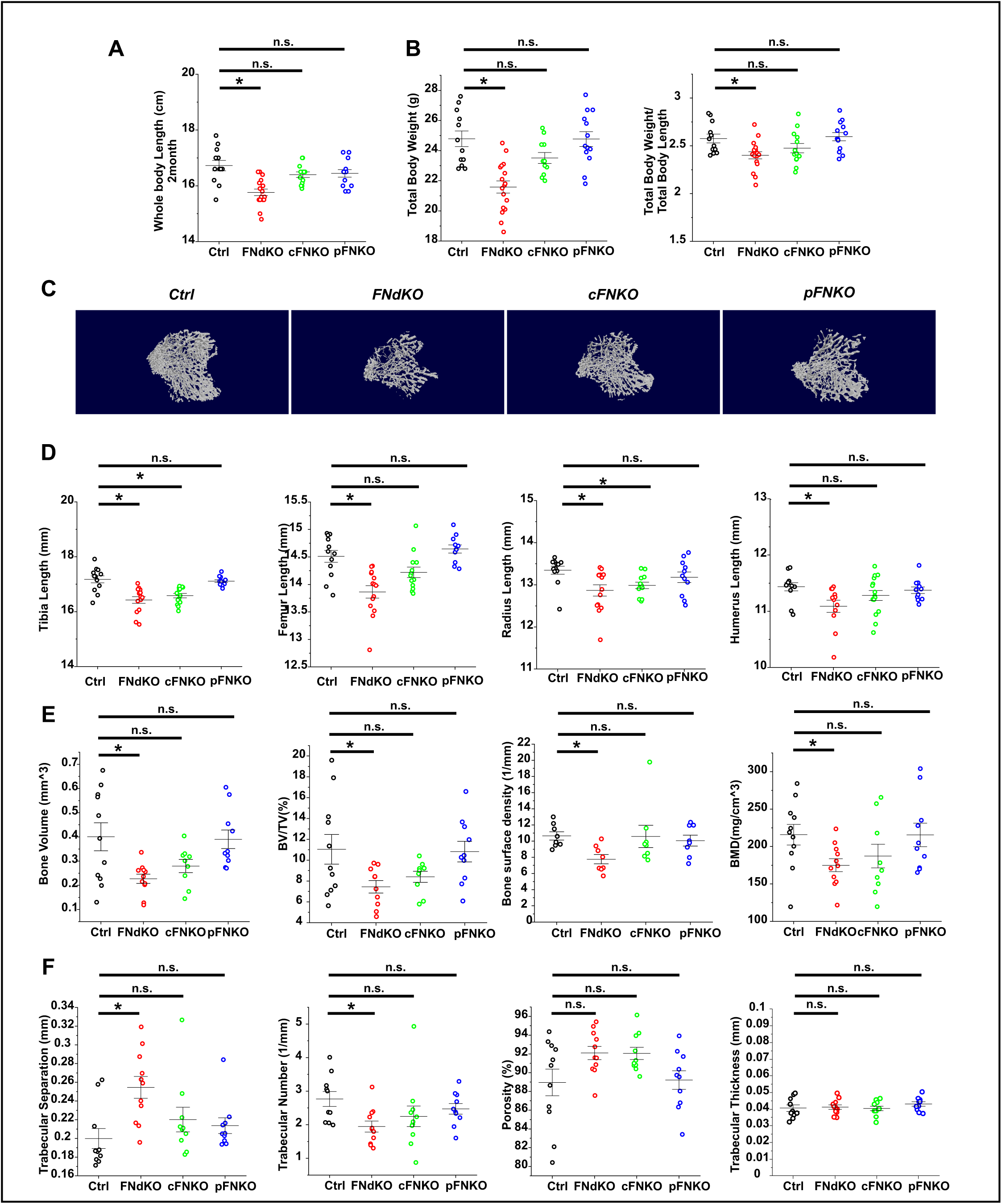
Morphological and morphometric analysis of adult control and FN knockout mice. **(A,B)** Measurements of whole-body length and total body weight normalized to total body length for 2-month-old control and FN knockout mice. Ctrl (n=12); FNdKO (n=17); cFNKO (n=11-13); pFNKO (n=13). **(C)** Representative micro-CT images of trabecular bone volume. **(D)** Quantification of the total bone length of the tibia, femur, radius, and humerus. Ctrl (n=12); FNdKO (n=12-14); cFNKO (n=12-15); pFNKO (n=11). (**E,F**) Quantification of trabecular bone parameters from micro-CT analyses. Ctrl (n=11); FNdKO (n=11); cFNKO (n=9); pFNKO (n=10). BV, Bone volume; TV, Tissue Volume; BMD, bone mineral density. Each data point represents one mouse. Error bars represent the standard error of the mean. * Represents a p-value of <0.05. “n.s.” indicates a non-significant p-value.

### Deletion of FN isoforms does not alter fibrillin-1 deposition in cartilage

Fibrillin-1 (FBN1) is a known regulator of bone development and growth ^41^, and previous studies have shown that an FN scaffold is essential for FBN1 assembly in cell culture ^42^ and *in vivo* in blood vessels ^33^. Hence, to investigate whether defects in FBN1 could potentially cause reduced bone growth in FNdKO mice, we immunostained P1 tibiae for FBN1 with specific antibodies not cross-reacting with FN. In control tibiae, FBN1 was prominently present around the growth plate in the perichondrium, in the resting chondrocyte zone within the growth plate, and in the trabecular bone **(Supp. Figure 6A)**. Surprisingly, no differences in FBN1 levels were qualitatively observed between the control and the single or double FN knockout samples **(Supp. Figure 6A)**. This was further validated by quantification of the total FBN1 levels within the growth plate **(Supp. Figure 6B)**. These results suggest that an FN network is not essential in the epiphyseal cartilage to facilitate FBN1 deposition and/or assembly.

### FN double-knockout mice are characterized by dysregulated chondrogenesis and bone mineralization

To analyze alterations in cartilage upon deletion of FN isoforms, we performed histological analysis of the growth plate. Safranin-O and Fast Green stained P1 tibia sections were used to determine the length of distinct chondrocyte zones **(Figure 5A)**. The FNdKO samples displayed a reduced length of the proliferating zone and an increase in the hypertrophic zone, whereas single FN knockout samples were not different from the control **(Figure 5B)**. This suggested dysregulated chondrogenesis in the FNdKO mice. Since FN promotes proteoglycan balance during chondrogenesis ^43^, we analyzed the proteoglycan levels by Alcian blue staining. No apparent changes in staining intensity were noted in the FN knockout samples **(Figure 5C)**. Immunostaining for the chondrogenic marker collagen type II (COLII) of P1 tibiae showed uniform COLII staining in both control and pFNKO samples **(Figure 6A**). However, the FNdKO and cFNKO samples displayed a disrupted uneven COLII deposition pattern in the growth plate **(Figure 6A).** Quantification showed significantly reduced total COLII protein in FNdKO samples but not in the cFNKO, which was further validated on the mRNA level (*Col2a1*) **(Figure 6B,C)**. Additionally, immunohistochemistry of the chondrocyte hypertrophy marker collagen type X (COLX) revealed increased COLX levels in the hypertrophic zone of FNdKO tibiae **(Figure 6D,E).** This was further confirmed by analysis of the *Col10a1* mRNA expression levels **(Figure 6F).**

**Figure 5.**
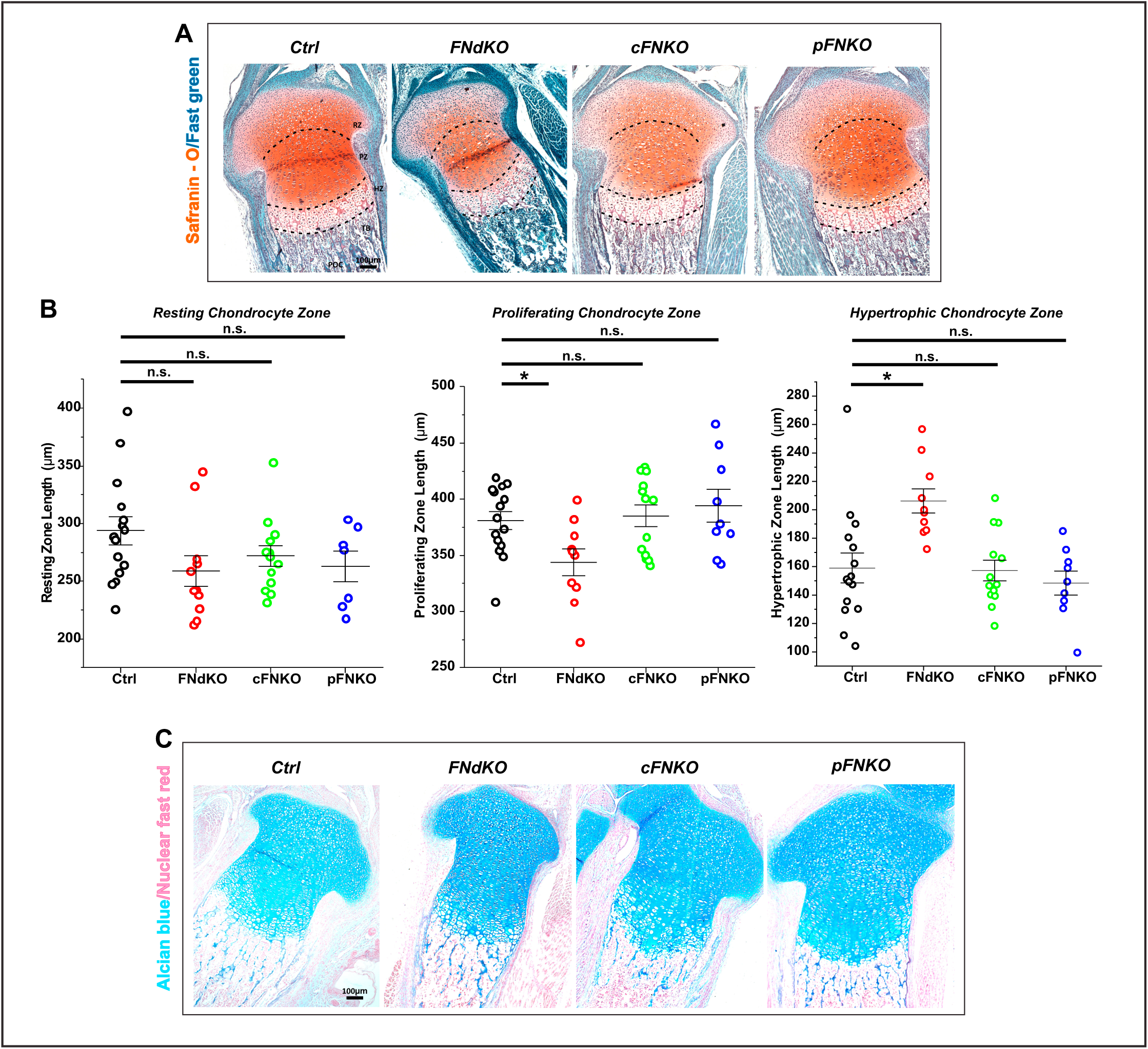
Histological analysis of the tibia growth plate of FN knockout mice. **(A)** Safranin O/Fast Green-stained sections of tibiae at P1 of control and FN knockout mice. Red dashed lines indicate the beginning/end of the resting chondrocyte zone (RZ), proliferating chondrocyte zone (PZ), hypertrophic zone (HZ), and trabecular bone (TB). **(B)** Quantification of the length of each chondrocyte zone. Ctrl (n=15); FNdKO (n=10-11); cFNKO (n=13); pFNKO (n=7-9). **(C)** Alcian blue (blue) stained sections of tibiae from P1 control and FN knockout mice. Ctrl (n=17); FNdKO (n=9); cFNKO (n=6); pFNKO (n=12). Each data point represents one mouse. Error bars represent the standard error of the mean. The scale bar represents 100 μm. * Represents a p-value of <0.05. “n.s.” indicates a non-significant p-value.

**Figure 6.**
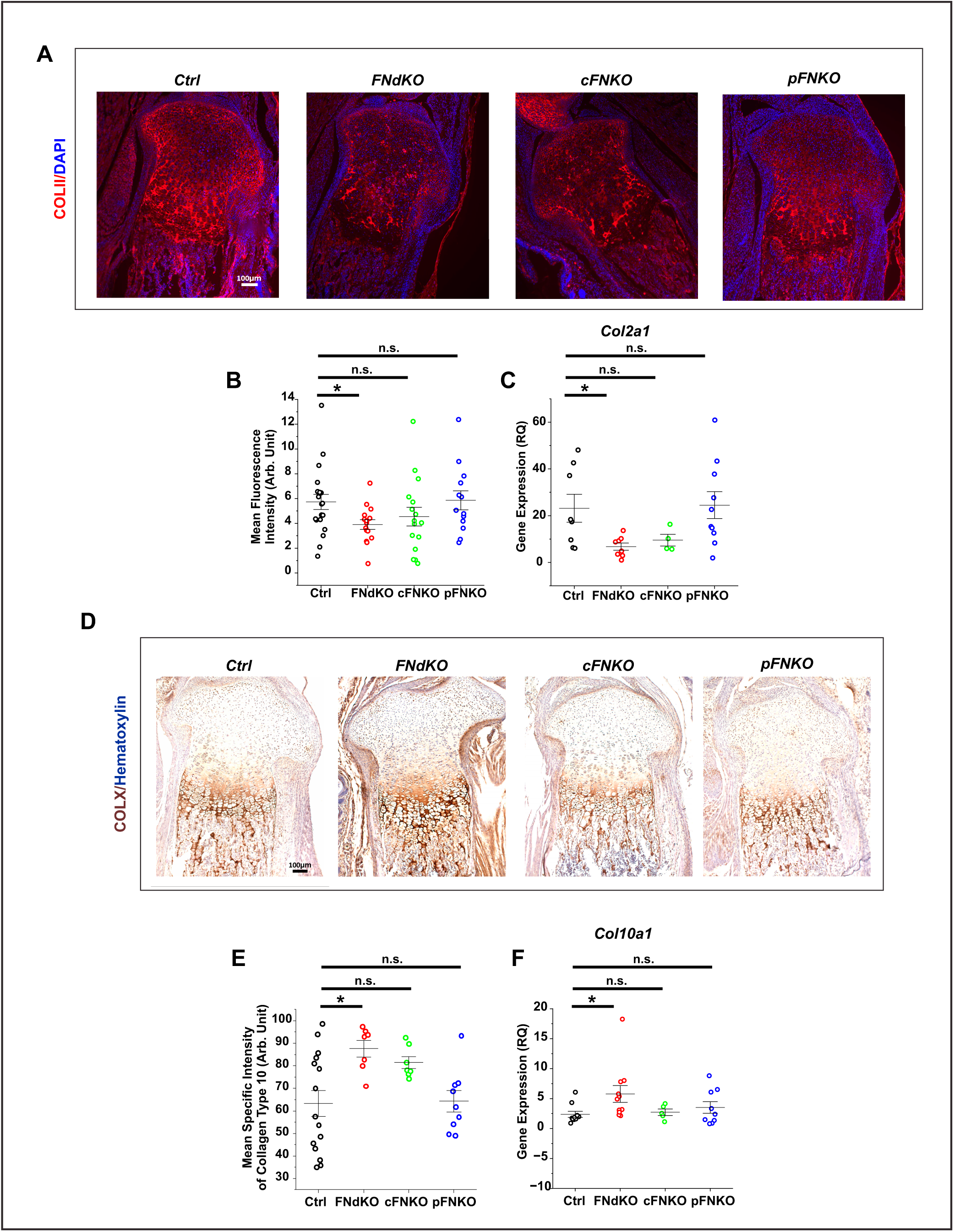
Dysregulation of chondrogenic markers in FN knockout mice. **(A)** Immunostaining of chondrogenic marker collagen type II (COLII, red) on tibia sections at P1 control and FN knockout mice. DAPI (blue) is used as a nuclear counterstain in all images. **(B)** Quantification of the mean fluorescence intensity of COLII in the growth plate. Ctrl (n=21); FNdKO (n=16); cFNKO (n=17); pFNKO (n=14). **(C)** Collagen type II gene (Col2a1) expression in the tibiae of control and FN knockout mice at postnatal day 1. Ctrl (n=8); FNdKO (n=8); cFNKO (n=4); pFNKO (n=10). **(D)** Immunohistochemistry of chondrogenic marker collagen type X (COLX, brown) on tibia sections of P1 control and FN knockout mice. Hematoxylin stain (purple) represents cell nuclei in all images. **(E)** Quantification of the mean specific intensity of COLX in P1 tibia sections. Ctrl (n=15); FNdKO (n=7); cFNKO (n=7); pFNKO (n=9). **(F)** Collagen type X mRNA (Col10a1) expression in the tibiae of control and FN knockout mice at P1. Ctrl (n=10); FNdKO (n=10); cFNKO (n=5); pFNKO (n=9). Each data point represents one mouse. Error bars represent the standard error of the mean. The scale bar represents 100 μm. * Represents a p-value of <0.05. “n.s.” indicates a non-significant p-value.

During endochondral ossification, the hypertrophic chondrocytes follow two fates: One where these cells terminally differentiate to initiate cartilage mineralization, providing a matrix for the osteoblasts, and then undergo apoptosis ^44^. The second fate is where they directly differentiate into osteoblasts and osteocytes ^6^. Hence, we analyzed these aspects in the FNKO models. Immunostaining for the cell apoptosis marker cleaved caspase-3 revealed significantly reduced apoptotic cells in the hypertrophic region of FNdKO tibiae at P1 but not of cFNKO and pFNKO **(Figure 7A,B)**. Consistent with these results, Von Kossa staining of tibiae demonstrated significantly diminished mineralized content in the trabecular bone only from the FNdKO mice **(Figure 7C,D)**. The FNdKO tibiae also had reduced mRNA steady-state levels for the bone maturation markers *Ssp1* and *Bglap* but not for *Sp7* and *Ibsp*; in contrast, the hypertrophy marker Sox9 was upregulated compared to controls (**Figure 7E**). None of the single FN knockout samples showed those changes. These data demonstrate that simultaneous deletion of the chondrocyte-specific cFN and pFN in developing cartilage of FNdKO alters chondrogenesis and the transition of cartilage to mineralized bone.

**Figure 7.**
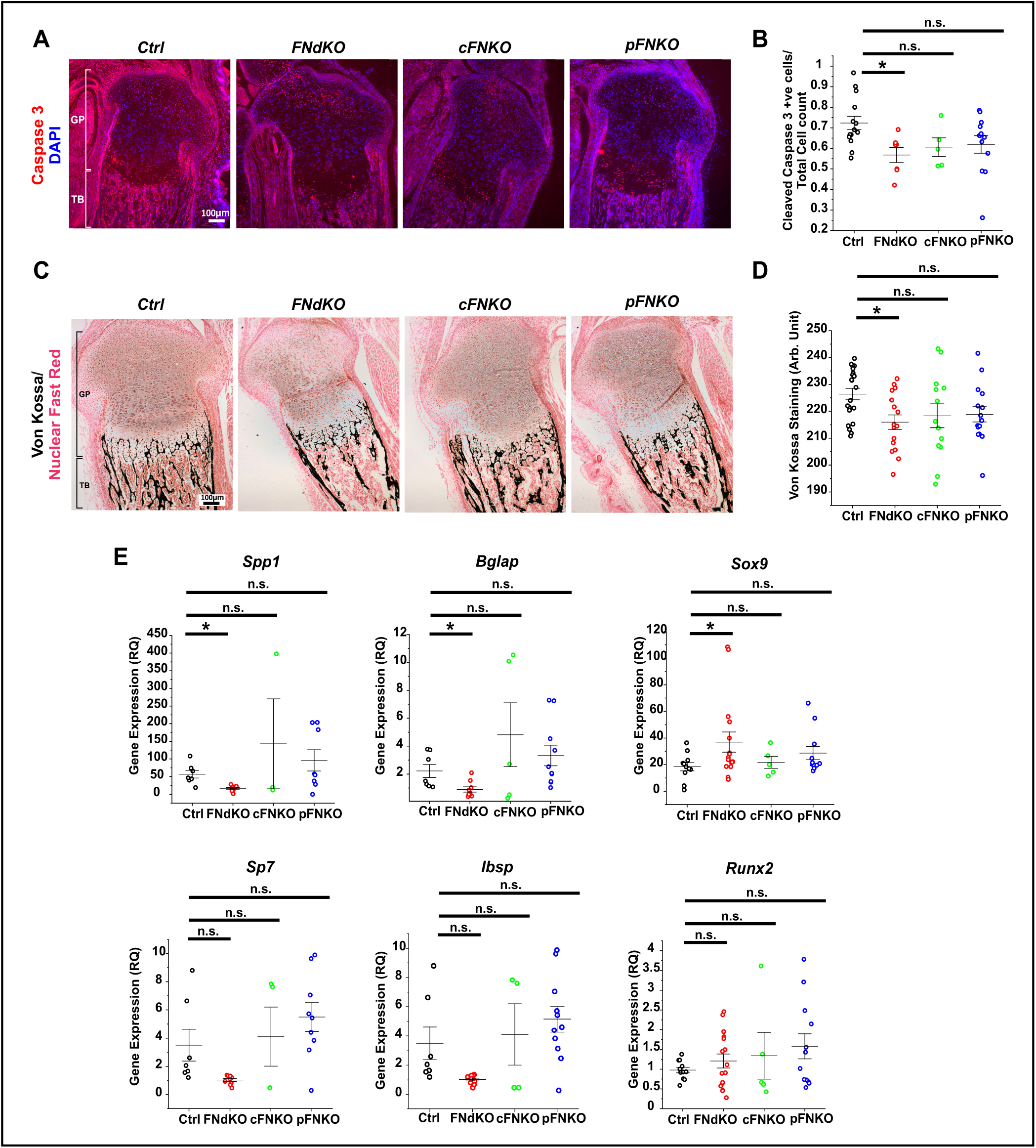
Altered development of trabecular bone in FN knockout mice. **(A)** Immunostaining for the cell apoptotic marker cleaved caspase 3 (red) in tibiae of P1 control and FN knockout mice. DAPI (blue) was used to counterstain the nuclei in all images. **(B)** Quantification of the caspase 3 positive cells normalized to total DAPI positive cells from images in **(A)**. Ctrl (n=14); FNdKO (n=7); cFNKO (n=5); pFNKO (n=12). **(C)** Von Kossa staining of calcium deposition (black) on tibia sections from P1 control and FN knockout mice. Note the mineralization in the trabecular bone of FNdKO mice. Nuclear fast red (pink) was used to counterstain cell nuclei in all images. **(D)** Quantification of total mineralized content from von Kossa-stained images in **(C)**. Ctrl (n=21); FNdKO (n=16); cFNKO (n=13); pFNKO (n=15). **(E)** Relative mRNA expression of chondrocyte hypertrophy (Sox9) and osteoblast markers (Spp1, Bglap, Sp7, Ibsp, and Runx2) in the tibiae of control and FN knockout mice at P1. Ctrl (n=7-11); FNdKO (n=9-16); cFNKO (n=3-5); pFNKO (n=8-12). Each data point represents one mouse. Error bars represent the standard error of the mean. The scale bar represents 100 μm. * Represents a p-value of <0.05. “n.s.” signifies a non-significant p-value.

### FN isoforms regulate chondrogenesis via TGFβ-mediated AKT signaling during postnatal bone development

TGFβ is an essential growth factor for regulated chondrogenesis ^45^, and TGFβ signaling regulates FN isoform levels during chondrogenesis and *vice versa* (refer to review ^34^). FN regulates TGFβ levels and its bioavailability via organizing LTBP1 assembly ^46^. It was also shown that FN regulates chondrogenesis via TGFβ-mediated PI3K/AKT signaling in a murine femur fracture healing model ^47^. Therefore, we investigated TGFβ levels and TGFβ-mediated AKT signaling in the FN knockout mice. Since TGFβ1 is the major TGFβ isoform present in bones ^48,49^, we tested the mRNA expression levels of TGFβ1 and its receptor TGFβR-1 in tibiae of P1 pups **(Figure 8A,B)**. FNdKO samples exhibited reduced levels of TGFβ1 concurrent with a significant increase in TGFβR-1 levels. The elevated TGFβ receptor mRNA may represent a compensatory response to the decreased mRNA levels of the ligand. Analysis of the downstream total and phospho-AKT levels by immunoblotting of protein lysates from P1 tibiae showed significant downregulation of phospho-AKT in samples from FNdKO but not from cFNKO and pFNKO mice **(Supp. Figure 7A-D)**. Immunostaining for phospho-AKT on P1 tibia sections showed reduced levels of active AKT in GP and in trabecular bone in FNdKO samples compared to controls **(Figure 8C,D)**. No significant changes were observed in single FN knockout tibiae. Overall, these findings demonstrate that the two FN isoforms together are essential to maintain TGFβ levels in cartilage, and deletion of both cFN and pFN results in reduced TGFβ-mediated AKT signaling in the developing cartilage.

**Figure 8.**
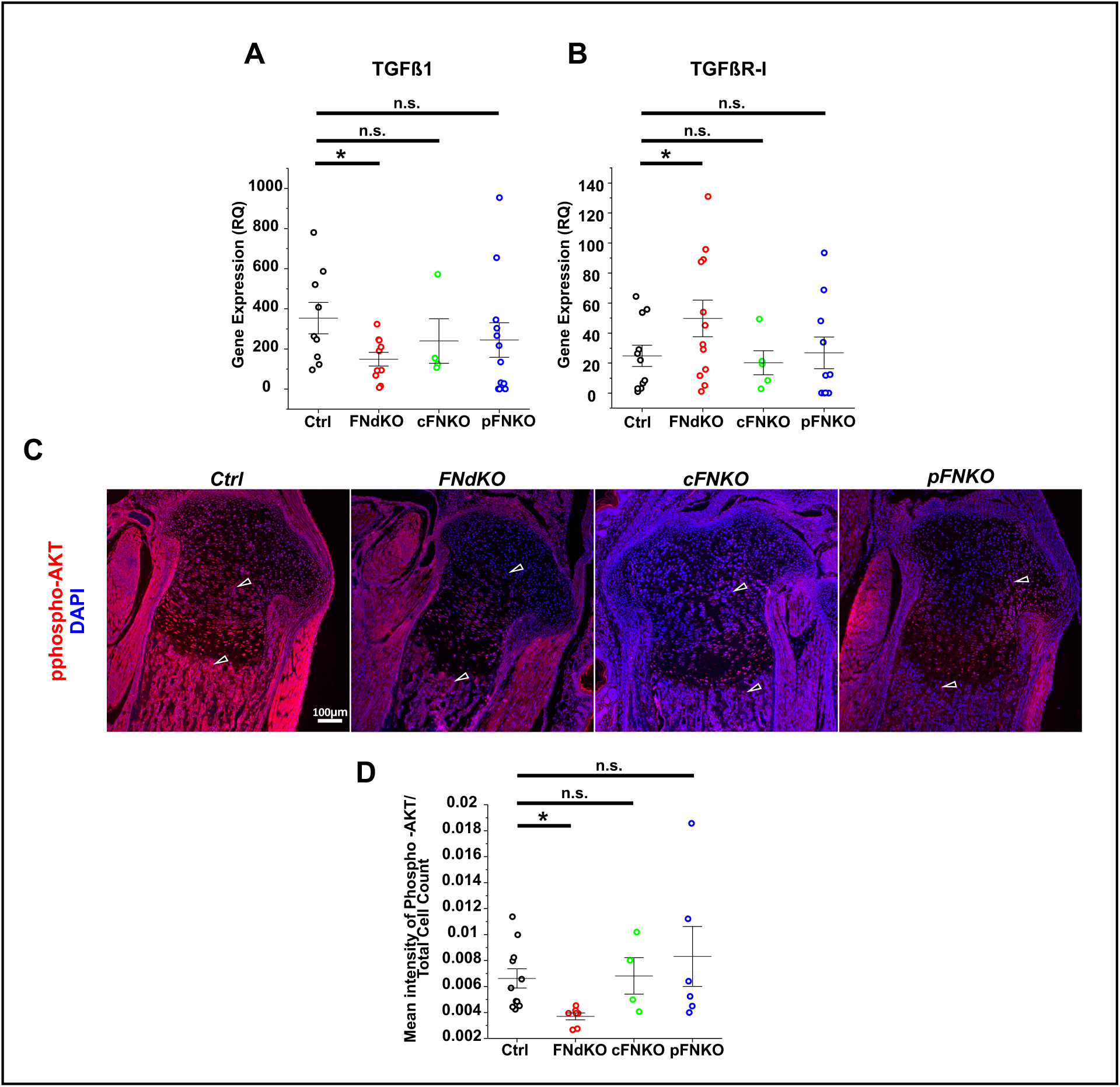
Analysis of TGFβ-mediated signaling in FN knockout mice. **(A-B)** Relative mRNA expression of TGFβ-1 and TGFβR-1 in the tibiae of control and FN knockout mice at P1. Ctrl (n=9-11); FNdKO (n=10-12); cFNKO (n=4-5); pFNKO (n=10-12). **(C)** Immunostaining for phospho-AKT (red) in tibiae obtained from P1 control and FN knockout mice. DAPI (blue) staining was used to visualize cell nuclei in all images. **(D)** Quantification of the mean fluorescence intensity of phospho-AKT, normalized to total DAPI positive cells from images in **(C)**. Ctrl (n=11); FNdKO (n=7); cFNKO (n=4); pFNKO (n=6). Each data point represents one mouse. Error bars represent the standard error of the mean. The scale bar represents 100 μm. * Represents a p-value of <0.05. “n.s.” marks a non-significant p-value.

## Discussion

FN is a promoter of chondrogenesis and essential for condensation of the stem cell mesenchyme, a prerequisite for chondrogenesis ^50^. However, the functional importance of FN during endochondral ossification and the roles of cartilage-specific cFN and the circulating pFN in this process are unknown and are thus the subject of this study.

Previous studies using cell culture models have demonstrated that FN is present during chondrogenesis ^51–54^. However, there is no evidence *in vivo* of how FN is expressed in limbs during endochondral ossification. In this study, we demonstrate that FN is present in mouse cartilage as early as E12.5 and is deposited in a unique pattern in the mouse limb with a distinct increase in specific growth plate zones (resting and hypertrophic) and the trabecular bone. pFN can enter several tissues and organ systems and compensate for the absence of cFN in vascularized tissues such as the aorta ^29,33^. Hence, we investigated this mechanism in the epiphyseal cartilage using cFNKO mice, which showed deletion of cFN as early as E16.5. In these mice, we expected that in the absence of cFN, pFN from the blood vasculature in the primary and secondary ossification centers would enter the growth plate to compensate for the lack of cFN. Surprisingly, at all analyzed fetal and neonatal time points, cFNKO mice showed FN deposition only in the trabecular bone and bone marrow but not in the growth plate. This contrasts with other tissues such as lungs, kidneys, heart, and blood vessels ^30^, where pFN can infiltrate from the vasculature. Based on these data, it is clear that cFN is the primary FN isoform in epiphyseal cartilage. In the growth plate, the transport of molecules and nutrients from adjacent blood vasculature occurs via diffusion ^55,56^. pFN is a large ∼440kDa dimer, and free diffusion from the blood vessel lumen is prevented by the vascular endothelial tight junctions ^56^. This was demonstrated in our previous work with fluorescently labeled dextran and with labeled pFN injected into mice, revealing that translocation of pFN did not occur via diffusion across the endothelial cell layer ^33^. Instead, a model of endothelial transcytosis was proposed that requires specific uptake of pFN and translocation through endothelial cells. The absence of pFN in the cFNKO growth plate suggests that such a compensatory mechanism does not exist in the cartilage, presumably due to the matrix composition of the growth plate. Immunostaining for total FN in the FNdKO showed that deletion of cFN in cartilage and circulating pFN leads to complete ablation of FN in the tibia. This was unexpected, as cells from other skeletal lineages, such as osteoblasts or the stromal mesenchymal cells, should synthesize and deposit cFN. This data suggests that the cFN synthesized in cartilage and pFN from blood translocating to the bone are the two major contributors of FN in a developing bone, not cFN produced from bone resident cells.

Previous studies have demonstrated that the expression of the transgene *Alb-Cre* in the Alb-(Cre/+) mouse model begins in the liver as early as E15.5 ^38^, and deletion of pFN was previously confirmed at P3 ^33,35^. In the present study, we demonstrate that pFN is significantly reduced in pFNKO mice at P1 and is completely deleted at 2 months. Further, *Bentmann et al.* previously reported that deletion of circulating FN in pFNKO mice causes a reduction in bone mineralization in adult mice ^35^. However, in the present study, we observed very little to no difference in bone mineralization for the pFNKO mice. It is possible that the sample size of 10 employed in our study was not sufficient to display these differences, as opposed to the very large sample size of 24-34 used by Bentmann *et al.* It is also possible that this discrepancy is potentially due to different mouse strains. Our mouse models were generated on a mixed background of C57BL/J6 and sv129, whereas the strain information in Bentmann *et al.* was not provided. For cFNKO mice, we observed a tendency for alterations in most of the analyzed parameters, but these did not reach statistical significance. It is possible that the absence of cFN in the growth plate induces subtle defects in chondrogenesis and trabecular bone formation early in development, which is rescued by the pFN once it enters the bone. On the contrary, FNdKO mice had complete ablation of cFN and circulating pFN in the developing bone, which explains the extent of differences in the phenotypes between the FNdKO and cFN mice.

Analysis of FNdKO mice at P1 showed a reduced length of all long bones, reduced ossification and shortening of vertebral bodies, irregular ossification of the growth plate and trabecular bone, dysregulation of chondrogenic markers (COLII and COLX), and reduced TGFβ1. These skeletal defects overlap with clinical phenotypes in SMDCF associated with FN mutations, leading to severe skeletal defects ^19–22^. However, some other pathological aspects are unique to SMDCF and were not observed in the conditional FN knockout mice. While the FNdKO mice showed a significant reduction in the length and ossification of the vertebrae, there was no indication of scoliosis, which is consistently present in SMDCF ^20–22^. Some SMDCF patients display pectus carinatum and defective mobility due to genu varum, but these phenotypes were absent in the conditional FN knockout mice. Additionally, some SMDCF patients exhibited craniofacial abnormalities with asymmetrical facial bone formation, which was not noticeable in the FN knockout mice. The most likely reason those skeletal aspects are unaffected by deletion of either cFN in chondrocytes or circulating pFN resides in the different ossification processes. The craniofacial bones develop mainly by intramembranous ossification involving direct differentiation of mesenchymal stem cells into osteoblasts, which express osteoblast-specific cFN that is not deleted in the conditional FN knockout mice used here ^35^. The FNdKO mice displayed a significant reduction of long bone length into adulthood, analyzed at 2 months. Adult FNdKO mice also exhibited reduced body length, bone microarchitecture, and bone mineral density, which overlaps with the clinical phenotypes of patients with SMDCF ^20–22^. Other parameters, such as the reduced body weight only developed in older mice (1- and 2-months), demonstrate a progressive increase in the severity of this phenotype with age. The reduction in body weight is presumably due to a decreased overall bone mineral density and shorter long bones that only become measurable as the long bones develop and grow.

Several cell culture studies position FN as an inhibitor of adipogenesis via integrin-mediated interactions that promote ERK/MAPK signaling ^57^. Additionally, assembled FN and its interaction with adipocytes via α5β1 integrin is essential for this anti-adipogenic activity, and disruption of FN assembly leads to increased adipogenesis in cell culture ^58^. We, therefore, analyzed the bone marrow adiposity in 2-month-old mice and identified that the adult FNdKO mice exhibit a significant increase in bone marrow adipocytes in the mature bone, which is concurrent with a reduction in BMD. Increased bone marrow adiposity indicates poor bone health associated with reduced BMD ^59–61^ and is associated with elevated bone fracture susceptibility in humans ^62,63^. Even though bone marrow adiposity was not analyzed in patients with SMDCF, affected individuals display reduced bone density and are prone to femoral fractures, suggesting a possible alteration in bone marrow fat. Moreover, recent studies have demonstrated that growth plate chondrocytes contribute to the bone marrow adipocyte population by transdifferentiation ^7^. Therefore, it is possible that the deletion of FN facilitates the differentiation of chondrocytes into adipocytes.

During chondrogenesis, both TGFβ1 and the downstream phospho-AKT signaling have essential roles in promoting chondrogenesis and are indispensable for endochondral ossification and skeletal development ^45,64–67^. TGFβ signaling promotes differentiation of MSCs into chondrocytes characterized by increased early chondrogenic markers such as collagen type II ^45,68,69^ and inhibition of chondrocyte hypertrophy associated with collagen type X reduction ^70^. AKT signaling also has similar functions of inhibiting chondrocyte hypertrophy ^66^. In the present study, TGFβ1 and phospho-AKT levels were significantly downregulated in the FNdKO tibia, demonstrating that simultaneous deletion of both cFN and pFN leads to altered TGFβ signaling. This reduction potentially contributes to the reduced collagen type II and increased chondrocyte hypertrophy markers *Col10* and *Sox9* in the FNdKO mice. Sox9 is an essential transcription factor that increases *Col10* gene expression in hypertrophic chondrocytes ^71^. Additionally, Sox9 prevents osteoblast differentiation by opposing the β-catenin pathway during chondrogenesis ^71^, suggesting that high levels of *Sox9* in FNdKO mice could prevent chondrocyte to osteoblast transdifferentiation.

FN is a chondrogenic and osteogenic agent that can facilitate the differentiation of MSCs into chondrocytes or osteoblasts ^39,72,73^. The combined loss of the cartilage-specific cFN and pFN in FNdKO bones led to reduced trabecular bone formation, reduced bone mineralization, and a significant reduction in the expression of bone-specific markers such as *Spp1* and *Bglap1*. Upon maturation in the growth plate, chondrocytes undergo cellular apoptosis or transdifferentiate into osteoblasts and osteocytes to form the trabecular bone ^6,8–10^. We observed that FNdKO mice exhibited reduced caspase 3 positive hypertrophic chondrocytes and reduced levels of specific bone markers. These findings suggest that the hypertrophic chondrocytes fail to undergo apoptosis and/or differentiation into osteoblasts in the absence of FN ^71^, leading to reduced bone formation and mineralization.

Figure 9 provides a working model based on the results of this study. In summary, we identified that FN is present early during embryonic limb development, following a unique deposition pattern during growth plate development and its transition into mature bone. The cellular FN is the major isoform in the growth plate, whereas pFN enters bone but not the growth plate, even in the absence of cartilage-specific cFN. This is different from the compensatory mechanism in other tissues. The combined ablation of cartilage-specific cFN and circulating pFN in FNdKO mice resulted in dysregulated chondrogenesis and trabecular bone formation, associated with reduced TGFβ1 and downstream phospho-AKT levels. These defects ultimately resulted in overall reduced bone growth and defective skeletal development. Single cFN and pFN knockout mice did not show these phenotypes. Our work establishes that cartilage-specific cFN and circulating pFN together are indispensable for regulated chondrogenesis, cartilage maturation, trabecular bone formation, and overall skeletal growth.

**Figure 9.**
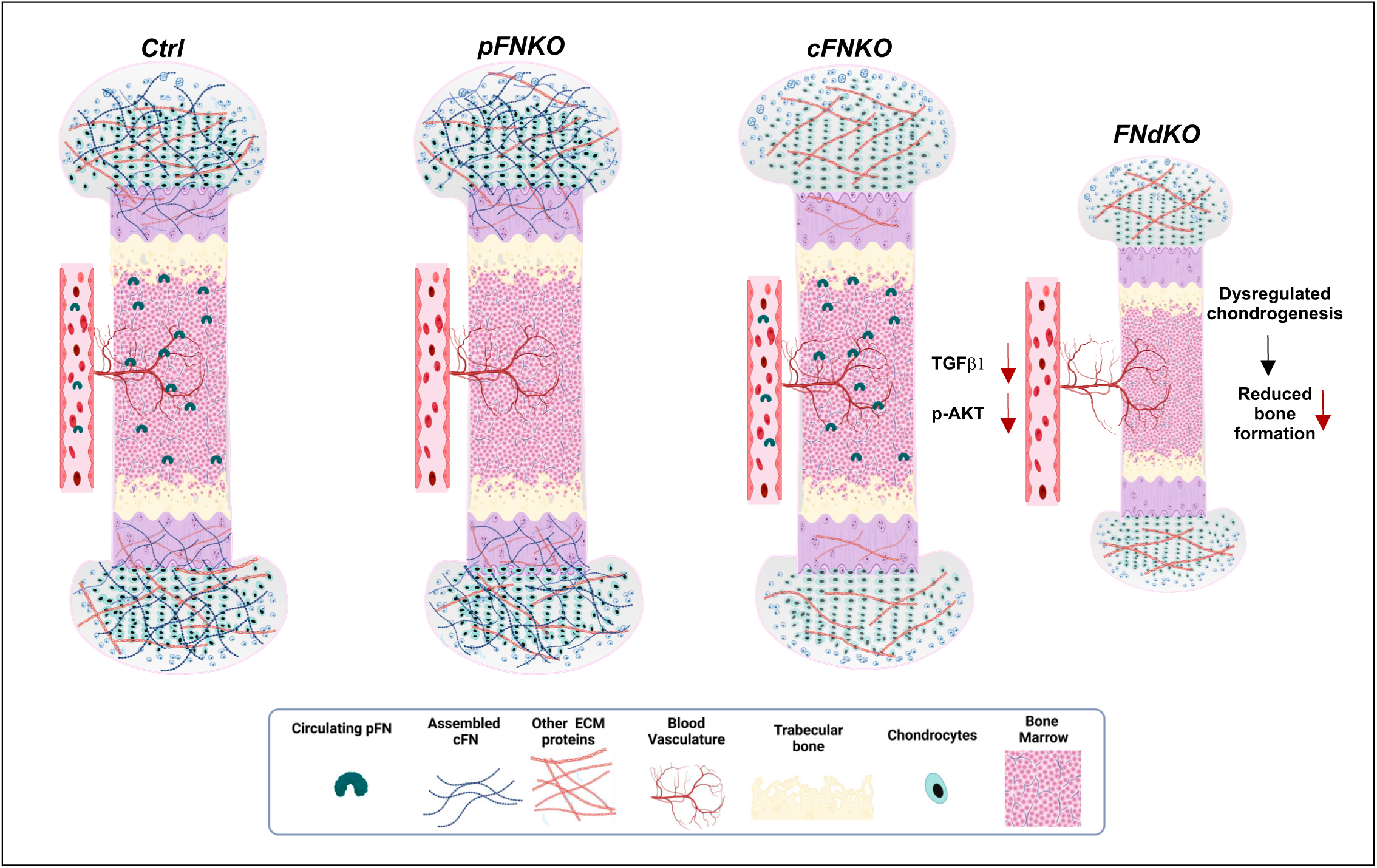
Working model of FN isoforms in cartilage and bone development. Schematic representation of the proposed model for the role of FN isoforms, cFN and pFN, in regulating chondrogenesis and bone growth. Details are discussed in the main text. The objects are not drawn to scale.

## Materials and Methods

### Ethics Statement

All animal experiments in this study were performed strictly following the guidelines established by the Canadian Council on Animal Care and were approved by the McGill University Animal Care Committee (protocol MCGL-5893).

### Mouse Models

FN*^flox/flox^* mice were received from Dr. Sarah Dallas, and the generation of this model is described in Sakai *et al.* ^32^. These mice were used as controls for all experiments. The Alb-Cre/+ (B6.Cg-Tg(Alb-cre)21Mgn/J; Stock No. 003574) mice were purchased from the Jackson Laboratory and were used to generate the liver-specific pFNKO mice by crossing the FN*^flox/flox^* with Alb-Cre/+ mice (FN*^flox/flox^*; Alb-Cre/+). The Col2a1-Cre/+ mice were crossed with FN*^flox/flox^* mice to generate the cartilage-specific cFNKO mice (FN*^flox/flox^*; Col2a1-Cre/+). The double knockout mice (FNdKO) were produced by crossing the two single knockout mice, resulting in the genotype FN*^flox/flox^*; Alb-Cre/+; Col2a1-Cre/+ (Figure 1C). Since the onset of puberty in mice occurs at P26 for females and at P30 for males ^74^, pups were not separated based on sex for experimental analyses at ages P1 to P15. Only male mice were employed for all studies performed at P30 and 2 months of age, as our initial analysis showed similar differences between ctrl and FNdKO for both male and female mice. All mice were housed under standard conditions (12 h of light-dark cycles) and were fed with facility chow. All mice were in a mixed background of C57BL/J6 and sv129. The sequences of primers used for genotyping of the mice are provided in Supplementary Table 2.

### Gross Morphological Analysis

Length measurements of bones from P1 old pups were performed after whole-mount skeletal staining (described below). Measuring calibers were used to determine the bone length of 1-month and 2-month-old mice. Body weight was determined using a standard weighing balance, and whole bone lengths were measured using a standard ruler.

### Whole-mount Skeletal Staining

P1 pups were euthanized and dissected to remove all internal organs and skin. The eviscerated skeletons were then stained with Alcian blue (cartilage) and Alizarin red (mineralized bone) using an established protocol ^75^. After 2-3 weeks required for complete clearance, the skeletons were imaged using an Amscope microscope (Model no. MU1000). The skeletons were further dissected to separate limbs, vertebrae, skull, and ribs, followed by imaging using a Stemi 2000-C microscope (Zeiss) and the AxioVision Rel.4.8 software (Zeiss). These images were used to quantify total and mineralized bone length with the ImageJ software ^76^.

### Micro Computed Tomography (Micro-CT)

Micro-CT was employed on 2-month-old tibia samples to determine the bone microarchitecture. All bone samples were scanned using a Skyscan 1272 microCT system (Bruker) at a resolution of 6 μm with 60-66kV voltage and 0.5 mm aluminum filter. The projection data was re-constructed using the NRecon software (Brucker). The trabecular bone region was selected starting from 100 μm below the end of the growth plate, extending up to 10% of the bone length. 3D morphometric parameters were determined using CT Analyzer software (Bruker) according to the guidelines provided by the American Society for Bone and Mineral Research ^77^. Trabecular bone parameters such as the total bone volume (BV in mm^3^), bone volume fraction (BV/TV in %), trabecular thickness (Tb.Th in mm), trabecular separation (Tb.Sp in mm), and trabecular number (Tb.N, mm^−1^) were determined. The bone mineral density (BMD) was determined from the mirror point values of each sample using the CTAn and MatLab software.

### Immunoblotting

For immunoblotting analysis, tibia samples of P1 pups were micro-dissected and sonicated at various experimental endpoints. Samples were lysed using RIPA buffer (50 mM Tris, pH 8.0, 150 mM NaCl, 0.5% sodium deoxycholate, 1% Triton X-100, and 0.1% sodium dodecyl sulfate) supplemented with 2% v/v protease inhibitor cocktail (Roche; Catalog #11697498001) and 1% v/v phosphatase inhibitor cocktail (Sigma-Aldrich; Catalog #P5726). Tissue lysates were centrifuged at 12,000×g for 5 min to pellet the debris and collect the protein extracts. Total protein concentrations of lysates were determined using the BCA protein assay kit (Thermo Fisher Scientific; Catalog #23225). 70µg of total protein was loaded and resolved on 6 or 10% SDS-PAGE gels, depending on the experimental setup. Gels were then wet-transferred to a 0.45 µm pore size nitrocellulose membrane (Bio-Rad; Catalog #1620115) and immunoblotted using specific primary antibodies, followed by incubation with *h*orse radish peroxidase-conjugated goat anti-rabbit or anti-mouse secondary antibodies (see **Supp. Table 1** for antibody details). Blots were developed using super signal chemiluminescent western blotting substrate (Thermo Fisher Scientific; Catalog# 34580) and imaged using the ChemiDoc MP system (Bio-Rad). Band intensities were quantified using ImageJ and normalized to the band intensities of the housekeeping protein.

### Quantitative PCR

Total RNA was extracted from micro-dissected tibia samples of P1 pups. Tissues were sonicated, and RNA was extracted using the RNeasy kit (Qiagen, Catalog #74104) per the manufacturer’s instructions. cDNA was prepared with 1 µg of total RNA using the Protoscript First Strand cDNA Synthesis kit (New England Biolabs; Catalog #E6560L) as instructed by the manufacturer. qPCR was performed using the SYBR Select Master Mix and a QuantStudio 5 Real-Time PCR System (Applied Biosystems). Primers against specific target genes are listed in **Supp. Table 2**. Gapdh was used as the control mRNA in all experiments. Relative quantification of mRNA levels (fold change) was calculated using the 2−ΔΔCt method.

### Tissue Staining and Microscopy

For all histological analyses, bone samples were harvested from mice at various embryonic and postnatal time points, followed by fixation in 4% paraformaldehyde for 24 h and embedding in paraffin. Bone samples from pups/mice of P15 to 2 months of age were decalcified after fixation with 14% w/v EDTA for 14 d. All paraffin-embedded samples were cut in sections of 5 μm thickness and used for histological and immunostaining procedures. Deparaffinized sections were stained with Hematoxylin and Eosin (general histology and bone marrow fat analysis), Safranin-O/Fast Green (cell orientation and growth plate length), and Von Kossa/Nuclear Fast Red (Mineralization) using standard procedures.

Levels and assembly of critical ECM molecules, such as total FN, fibrillin-1 (FBN1), collagen type II, as well as several intracellular proteins, were determined using indirect immunofluorescence staining. Depending on the marker analyzed, the sections were treated for antigen retrieval with 10 mM citric acid, 0.05% Tween 20 (pH 6) at 98°C and/or treatment with bacterial type XXIV proteinase (10 μg/mL; Sigma; Catalog #P8038). The sections were blocked with 2% bovine serum albumin (VWR; Catalog #10842-692), incubated with primary antibodies at specific dilutions (**Supp. Table 1**), followed by staining with the respective fluorophore-conjugated goat anti-rabbit or anti-mouse secondary antibodies (**Supp. Table 1**). An aqueous mounting medium including 4, 6-diamidino-2-phenylindole (DAPI) (Abcam; Catalog #188804) was used to counterstain cell nuclei. Immunofluorescence imaging was performed using an epifluorescence microscope AxioImager M2 (Zeiss) with an Orca Flash 4.0 CMOS grayscale camera. All images were originally in grayscale and then pseudocolored using the Zen Pro software version 2.6 (Zeiss). All brightfield imaging was performed using an ICC5 camera and the AxioImager M2. Histological and immunostained sample images were analyzed and quantified using ImageJ as described under the *Image Quantifications* section.

### Immunohistochemistry (IHC)

For IHC analysis, tissue sections were deparaffinized and rehydrated as described in the previous methods section. Sections were treated for antigen retrieval with 10 mM citric acid, 0.05% Tween 20 (pH 6) at 98°C, followed by treatment with bacterial type XXIV proteinase. For analysis of collagen type X, sections were additionally treated with 5mg/mL hyaluronidase (Sigma; Catalog #H3506) in PBS for 1 h at 37°C. The sections were blocked with 2% bovine serum albumin and incubated with primary antibody as described above. Samples were further treated with the HRP conjugated mouse polymer and developed using the Dako Envision+ System-HRP (DAB) staining system (Catalog #K400111-2, #K346811-2) following the procedure provided by the manufacturer. Samples were imaged using the ICC5 camera and AxioImager M2. Images were analyzed and quantified using the ImageJ software as described under the *Image Quantifications* section.

### Image Quantifications

Quantification of the mean fluorescence intensities of specific markers was performed using ImageJ. Cleaved caspase 3 positive cells were quantified by outlining the hypertrophic growth plate zone using ImageJ and manually counting the total cells and stained marker-positive cells. Quantification for intracellular marker phospho-AKT was performed by determining the mean intensity of phospho-AKT in the growth plate and bone region, followed by normalization to the total cell count. Quantification of nuclei was conducted as described previously ^78^. All marker-positive cells were normalized to the total number of cells per image. Length measurements of growth plate zones were performed using the line tool in ImageJ. For all image analyses, the scale was set based on the scale bar per image of the data set.

### Statistics

All data are represented as means ± Standard Deviation (SD) or Standard Error of the Mean (SEM) depending on the experimental setup as indicated in the figure legends. Significance was evaluated using Student’s two-tailed t-test. All statistical analyses were performed using the OriginPro version 2023 software (OriginLab). Outlier analysis was conducted using the Grubb’s test with confidence levels of 95%. Data with p ≤ 0.05 were considered significant and are denoted as *. Non-significance is labeled “n.s.”.

## Schematic Figures

All schematics in this manuscript were prepared using the Biorender software (https://www.biorender.com).

## Data availability

The data sets of this study are available from the corresponding author upon request.

## Acknowledgments

We express our sincere gratitude to Dr. Sarah Dallas and Dr. Takao Sakai for providing the FN^flox/flox^ mice, Dr. Rene St-Arnaud for providing the Col2a1-Cre/+ mice, and Ms. Olivia Vaikla for assisting in the quantifications of the whole-mount skeletal staining, and Ms. Melanie Dormann for assisting in mouse dissections and histological analyses.

## Author Contributions

ND, PC, and DPR conceived the study, designed the experiments, analyzed the data, and wrote the manuscript (with critical input from all authors). ND performed the experiments. PC and DPR provided the research funds.

## Competing Interests

The authors declare no competing interests.

## Funding

This work was supported by the Canadian Institutes of Health Research (Grant PJT-156140 to PC and DPR), Réseau de recherche en santé buccodentaire et Osseuse (PC and DPR) and the Fonds de recherche de Quebec (Dossier No. 291220, fellowship to ND). The funders have no role in study design, data collection and analysis, decision to publish, or manuscript preparation.

## Notes

### Competing Interest Statement

The authors have declared no competing interest.

