## Supplementary Information for "Role of Fibronectin in Postnatal Skeletal Development"

**Running Title:** Fibronectin in skeletal development

**Keywords:** Fibronectin, Skeletal Development, Chondrogenesis, Differentiation, Chondrocytes, Growth Plate

#### Supplementary Methods

##### *Analysis of Mouse Plasma Samples*

To analyze plasma samples from mice, blood was collected at various experimental time points by intracardiac puncture after administering an anesthetic cocktail. Plasma was prepared from blood using EDTA-coated tubes (Sarstedt) by centrifuging blood samples at 8,000×g for 8 min at 4°C. 0.1 µL of plasma per sample was analyzed for FN by standard western blotting using 6% SDS gels under reducing conditions (5% 2-mercaptoethanol). Polyclonal rabbit anti-mouse FN antiserum was employed as the primary antibody, and Peroxidase-AffiniPure Goat Anti-Rabbit IgG (H+L) was used as the secondary antibody **(Supp. Table 1)**. Immunoblots were developed as described above under the section for Immunoblotting.

##### *Bone Marrow Adipocyte Quantification*

Alterations in bone marrow fat were determined through histological H&E staining of the bone sections. Stained sections were analyzed by brightfield microscopy, and adipocytes were counted on H&E-stained images. For all image analyses, the scale was set based on the scale bar per image of the data set.

### Supplementary Figure 1

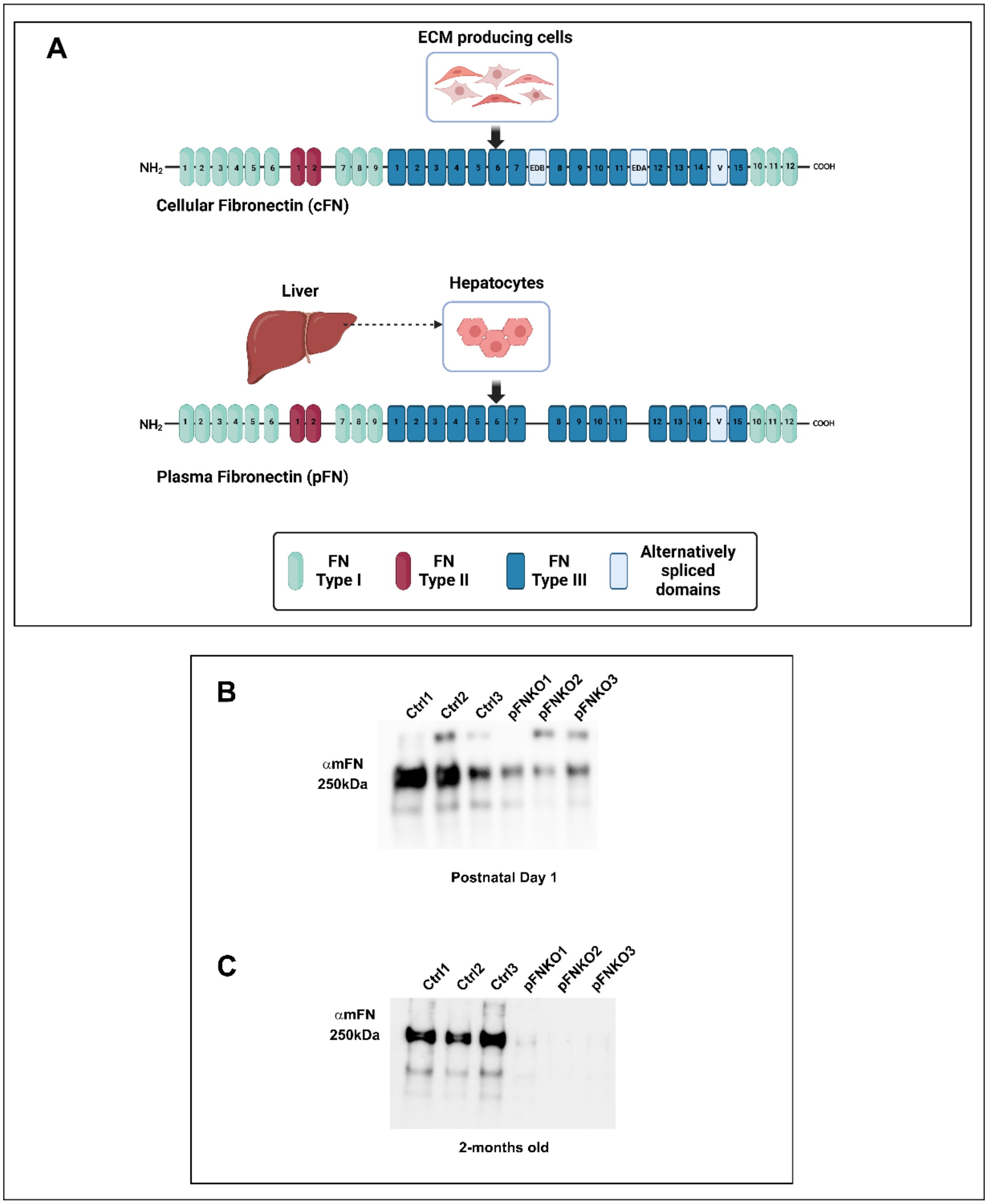


**Supp. Figure 1. Overview of FN isoforms and deletion of pFN in the FN knockout mice.**

**(A)** Schematic representation of the two principal isoforms of FN and their physiological sources. Note that the exons coding for EDA, EDB, and V domains can be alternatively spliced to produce FN isoforms with various combinations of these domains. **(B-C)** Immunoblotting for total FN (~250 kDa) in serum from control (n=3) and pFNKO mice (n=3) at P1 and 2 months of age. Samples were separated on 6% SDS gels under reducing conditions with 5% 2-mercaptoethanol. Relative to the control, decreasing levels of FN were observed in pFNKO at P1 age and complete absence at 2 months of age.

#### Supplementary Figure 2

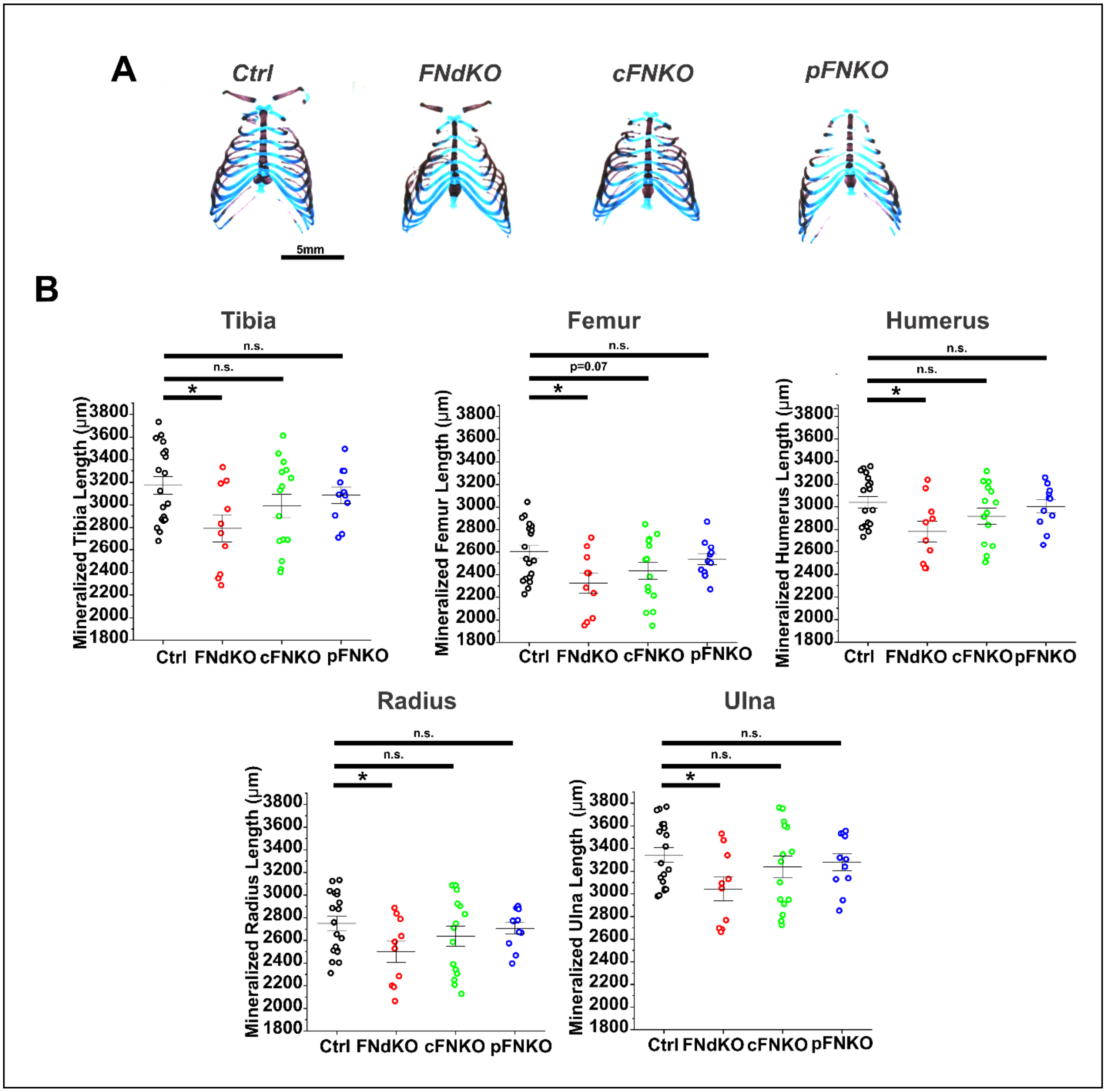


**Supp. Figure 2. Analysis of rib cages and mineralized bone length of P1 old mouse pups.**

**(A)** Representative images of P1 old mouse ribcages using whole-mount skeletal staining with alcian blue (blue, cartilage) and alizarin red (red, mineralized bone). The scale bar represents 5 mm. **(B)** Quantification of the mineralized bone length of the tibia, femur, humerus, radius, and ulna. Each data point represents one mouse pup. Error bars represent the standard error of the mean. Ctrl (n=19); FNdKO (n=10); cFNKO (n=15); pFNKO (n=11). * Represents a p-value of <0.05. "n.s." indicates non-significant p-values.

### Supplementary Figure 3

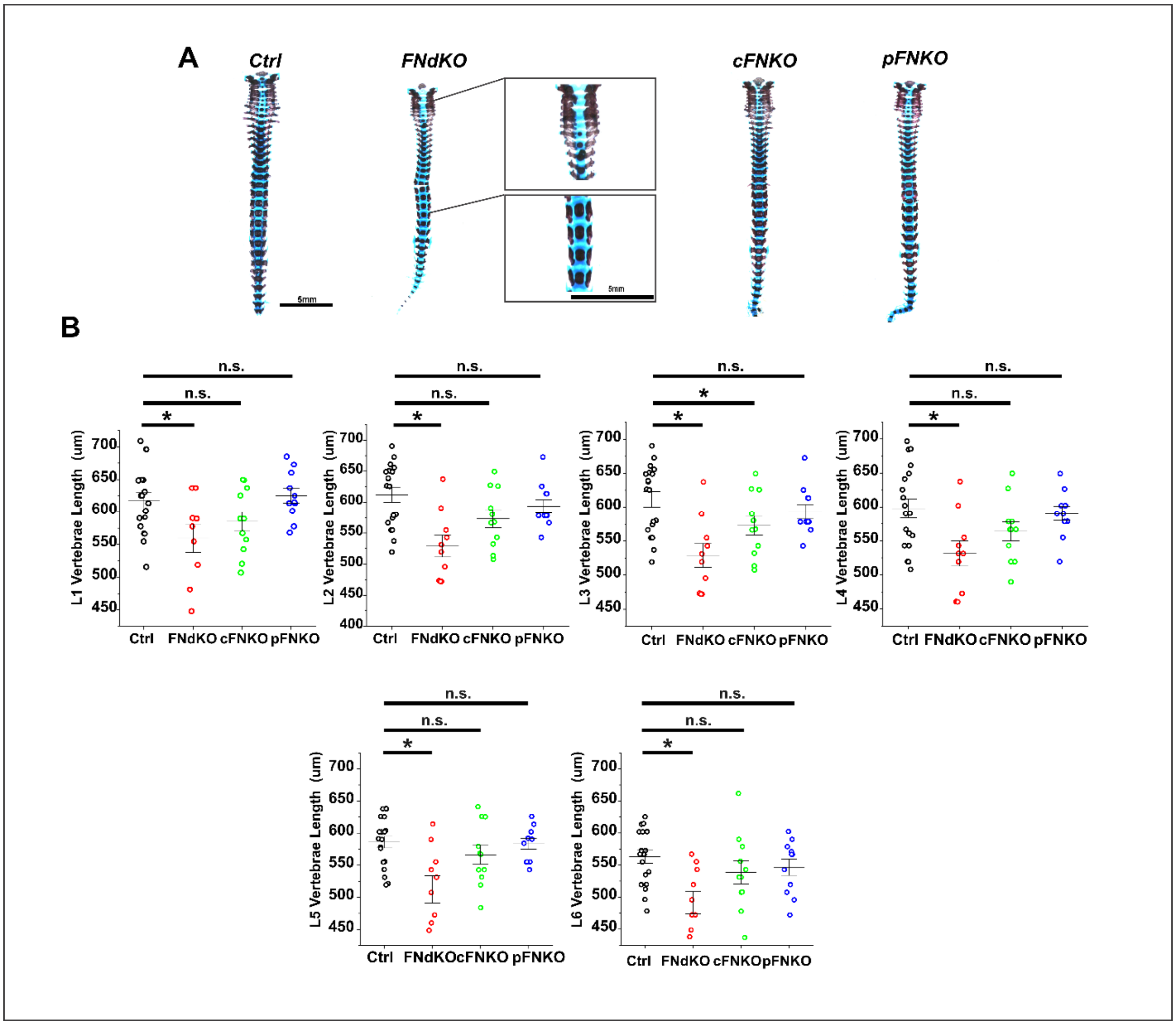


**Supp. Figure 3. Assessment of abnormalities in the vertebral column of P1 old mouse pups.**

**(A)** Representative images of P1 old mouse vertebral columns using whole mount skeletal staining with alcian blue (blue, cartilage) and alizarin red (red, mineralized bone). Note the reduced number of mineralized nodules in the upper vertebral column of FNdKO and the overall reduction in the region of ossification in all vertebral bodies. The scale bar represents 5 mm. **(B)** Quantification of the length of lumbar vertebrae from L1 to L6. Each data point represents one pup. Error bars represent the standard error of the mean. Ctrl (n=19); FNdKO (n=10); cFNKO (n=11); pFNKO (n=11). * Represents a p-value of <0.05. "n.s" indicates non-significant p-values.

### Supplementary Figure 4

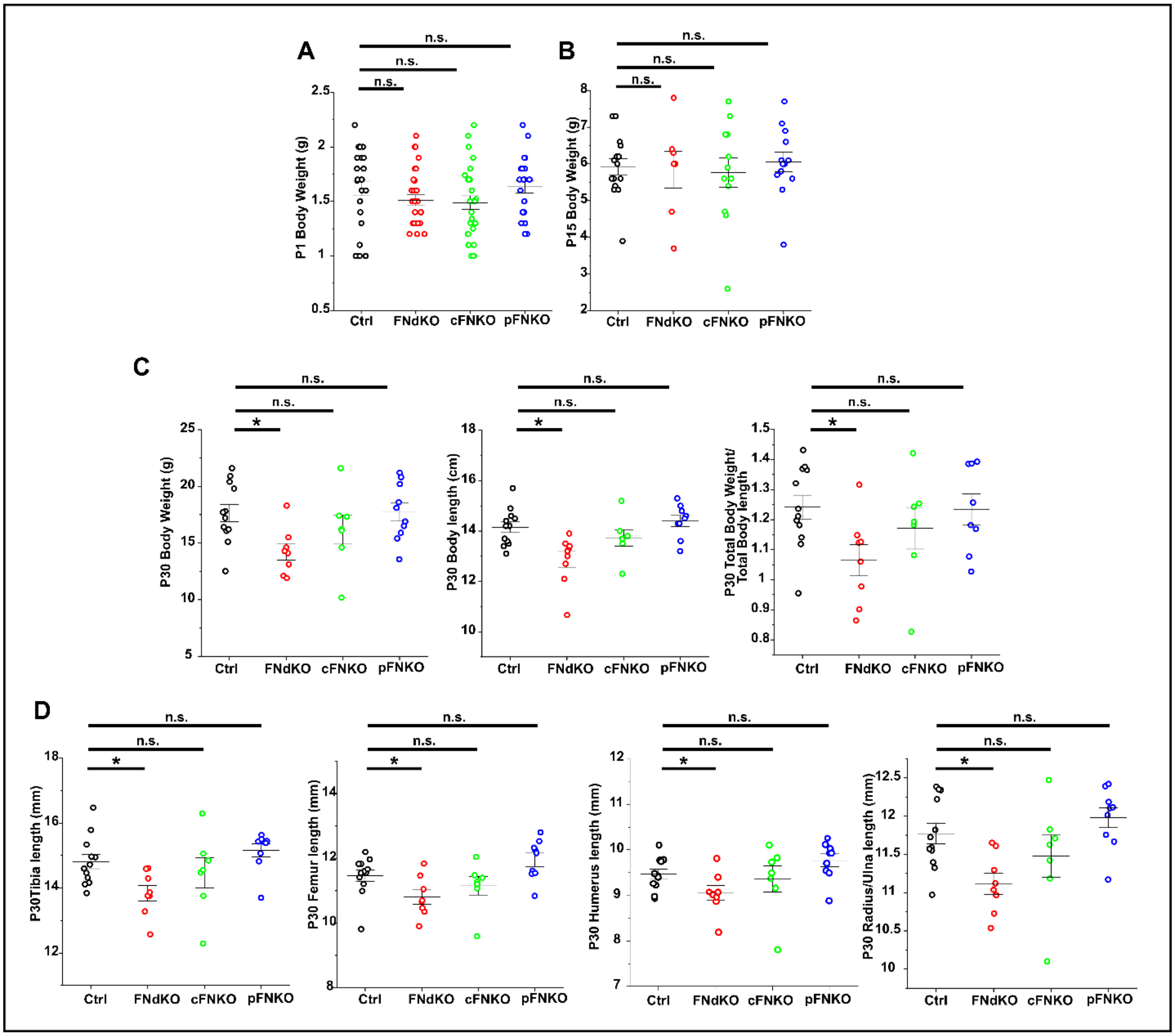


**Supp. Figure 4. Gross morphological analysis of the FN knockout mice at P15 and 1 month of age.**

**(A)** Quantification of total body weight of P1 old pups. Ctrl (n=27); FNdKO (n=29); cFNKO (n=29); pFNKO (n=23). **(B)** Quantification of the total body weight of P15 old pups. Ctrl (n=15); FNdKO (n=7); cFNKO (n=12); pFNKO (n=13). **(C)** Measurement of body weight, body length, and total body weight normalized to body length of P30 old mice. Ctrl (n=12); FNdKO (n=8-9); cFNKO (n=7); pFNKO (n=8-10). **(D)** Quantification of the total bone length of the tibia, femur, humerus, and radius of P30 old mice. Ctrl (n=11-12); FNdKO (n=8-9); cFNKO (n=7); pFNKO (n=9). Each data point represents one pup or mouse. Error bars represent the standard error of the mean. * Represents a p-value of <0. “n.s.” indicates non-significant p-values.

### Supplementary Figure 5

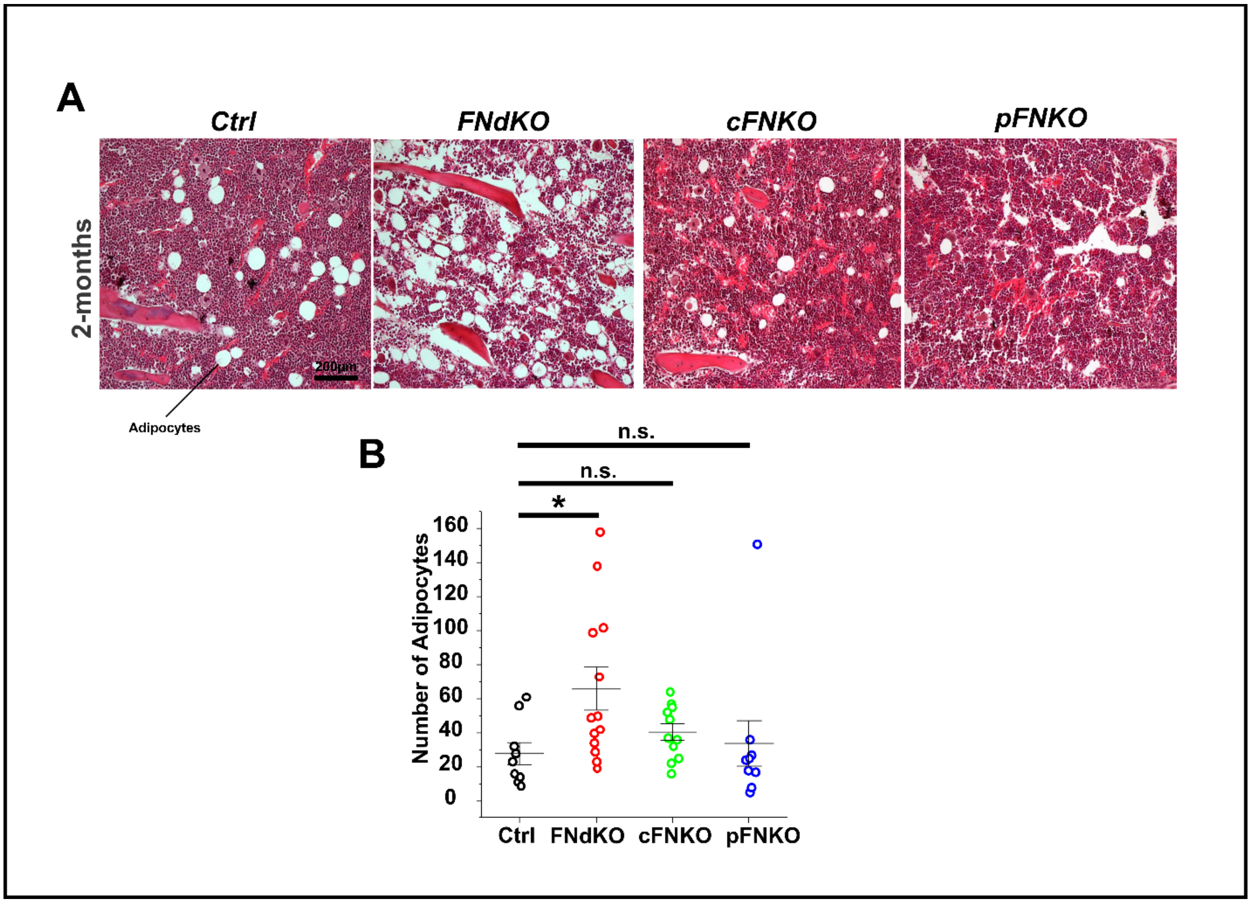


**Supp. Figure 5. Bone marrow fat analysis of the FN knockout mice.**

**(A)** Hematoxylin (purple) and eosin (pink) (H&E) stained images of bone marrow from tibiae of 2-month-old male control and FN knockout mice as indicated. Note the increased number of adipocytes in the bone marrow of FNdKO mice. Hematoxylin staining marks the cell nuclei in all images. The scale bar represents 200 µm. **(B)** Quantification of adipocyte numbers from H&E-stained images in **(A)**. Ctrl (n=9); FNdKO (n=13); cFNKO (n=11); pFNKO (n=10). Each data point represents one mouse. Error bars represent the standard error of the mean. * Represents a p-value of <0.05. "n.s." indicates non-significant p-values.

### Supplementary Figure 6

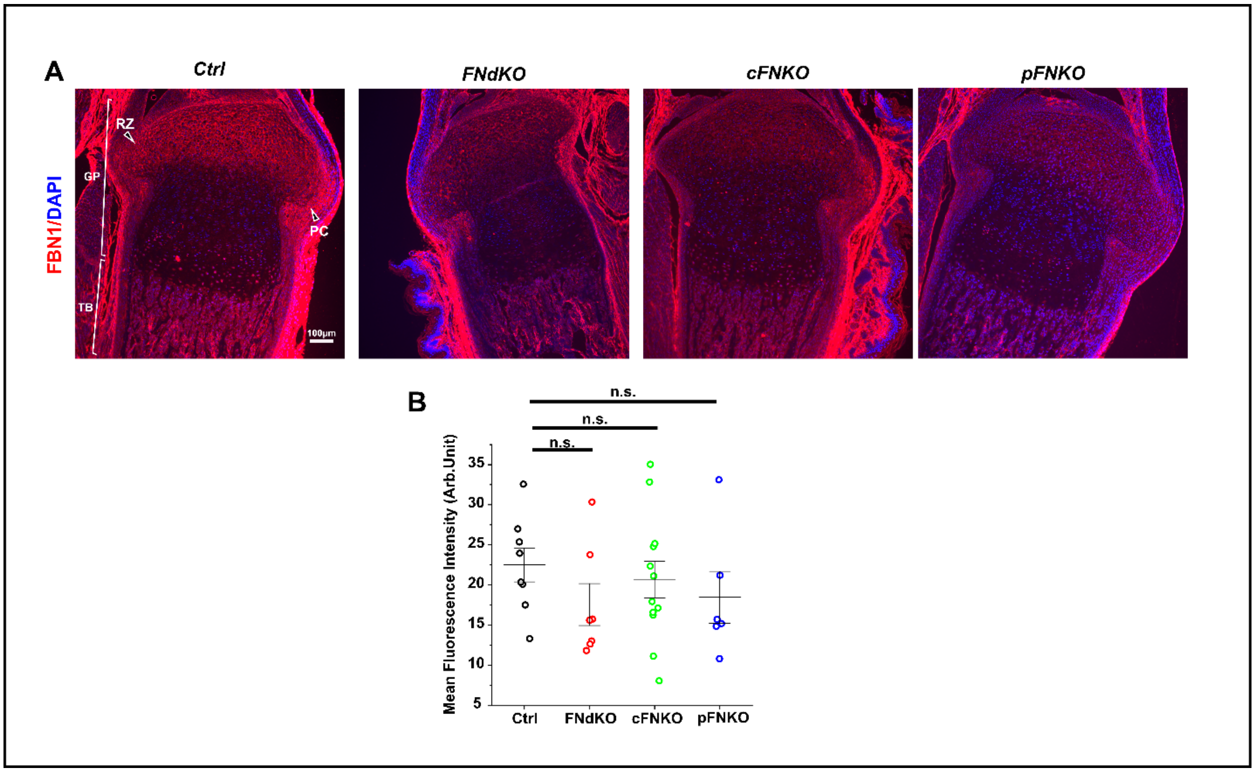


**Supp. Figure 6. Fibrillin-1 is not altered in the growth plate of FN knockout mice.**

**(A)** Immunostaining for fibrillin-1 (FBN1) deposition (red) in tibia sections from P1 old control and FN knockout mice. In all samples, note the distinct FBN1 staining in the growth plate's resting chondrocyte zone (RZ). DAPI staining (blue) represents the cell nuclei in all images. The scale bar represents 100 μm. **(B)** Quantification of mean fluorescence intensity of FBN1 from the growth plate region of the images shown in **(A).** Ctrl (n=8); FNdKO (n=7); cFNKO (n=12); pFNKO (n=6). Each data point represents one mouse. Error bars represent the standard error of the mean. * Represents a p-value of <0.05. "n.s." indicates non-significant p-values. Growth plate, GP; Trabecular bone, TB; Resting chondrocyte zone, RZ; Perichondrium, PC.

### Supplementary Figure 7

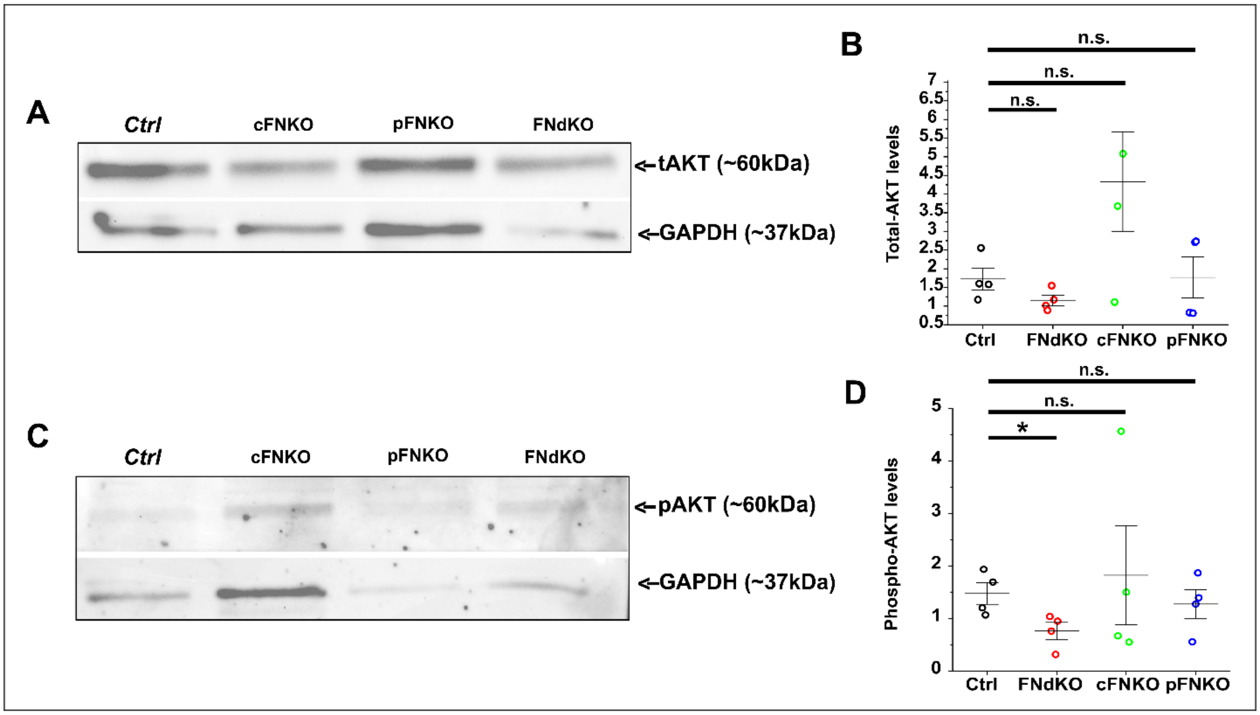


**Supp. Figure 7. Analysis of total and phospho-AKT levels in tibiae of FN knockout mice.**

**(A,B)** Immunoblotting of total AKT (tAKT, 60 kDa) and GAPDH (37 kDa) using protein lysates from P1 old control and FN knockout tibiae. A representative immunoblot is shown in **(A)**, and densitometric analysis of tAKT normalized to GAPDH is displayed in B (n=4 per group). **(C,D)** Immunoblotting of phospho-AKT (pAKT, 60 kDa) and GAPDH (37 kDa) using protein lysates from P1 old control and FN knockout tibiae. A representative immunoblot is shown in **(C)**, and densitometric analysis of pAKT normalized to GAPDH is displayed in **(D)**. * Represents a p-value of <0.05. "n.s." indicates non-significant p-values.

**Supplementary Table 1.** List of Antibodies used in this study.

| *List of Antibodies* | *Catalog Number* | *Source* | *Experimental Use (Dilution)* |
|---|---|---|---|
| Anti-Mouse FN antibody | AB2033 | Millipore Sigma | Immunoblotting (1:1000) and Immunostaining (1:500) |
| Anti-collagen type 2 antibody | II-II6B3 | DSHB | Immunostaining (1:20) |
| Anti-GAPDH antibody | 2118S | Cell Signaling | Immunoblotting (1:1000) |
| Anti-collagen type X antibody | 14-9771-82 | Invitrogen | Immunohistochemistry (1:200) |
| Anti-Cleaved Caspase 3 antibody | 9664P | Cell Signaling | Immunostaining (1:500) |
| Anti-Phospho-AKT antibody | AB38449 | Abcam | Immunoblotting (1:500) and Immunostaining (1:200) |
| Anti-Total AKT antibody | 9272S | Cell Signaling | Immunoblotting (1:500) |
| Anti-Fibrillin-1 antibody | - | Lab-generated* | Immunostaining (1:500) |
| Anti-Phospho-ERK1/2 antibody | 4377S | Cell Signaling | Immunostaining (1:200) |
| Goat anti-Mouse IgG (H+L) Cross-Adsorbed Secondary Antibody, Cyanine5 | A10524 | Invitrogen | Immunostaining (1:200) |
| Goat anti-Mouse IgG (H+L) Cross-Adsorbed Secondary Antibody, Cyanine3 | A10521 | Invitrogen | Immunostaining (1:200) |
| Peroxidase-Conjugated AffiniPure Goat Anti-Mouse IgG (H+L) | 115-035-003 | Jackson ImmunoResearch | Immunoblotting (1:500) |
| Peroxidase-Conjugated AffiniPure Goat Anti-Rabbit IgG (H+L) | 111-035-003 | Jackson ImmunoResearch | Immunoblotting (1:500) |

* Shi Y, Jones W, Beatty W, Tan Q, Mecham R, Kumra H, Reinhardt DP, Gibson MA, Reilly MA, Rodriguez J, Bassnett S. (2021). Matrix Biol 95, 15-31.

**Supplementary Table 2.** Primer sequences used in this study.

| *Gene* | *Strand* | *Sequence 5'-3'* | *Experimental Use* |
|---|---|---|---|
| ***FN floxed transgene*** | Sense | GTACTGTCCCATATAAGCCTCTG | Genotyping |
| | Antisense | CTGAGCATCTTGAGTGGATGGGA | Genotyping |
| ***Col2a1-Cre transgene*** | Sense | CAGCAGAACTCCGAGGAAAG | Genotyping |
| | Antisense | CATCGACCGGTAATGCAGG | Genotyping |
| | Sense | CTAGGCCACAGAATTGAAAGATCT | Genotyping |
| | Antisense | GTAGGTGGAAATTCTAGCATCATCC | Genotyping |
| ***Alb-Cre transgene*** | Sense (Alb) | CCTGCCAGCCATGGATATAA | Genotyping |
| | Antisense (Alb) | GTTGTCCTTTGTGCTGCTGA | Genotyping |
| | Antisense (transgene) | GAAGCAGAAGCTTAGGAAGATGG | Genotyping |
| ***Gapdh*** | Sense | GTTGCCATCAACGACCCCTTC | Real-time qPCR |
| | Antisense | GTTGCCATCAACGACCCCTTC | Real-time qPCR |
| ***Sp7*** | Sense | TGCCAGTAATCTTCAAGCCAG | Real-time qPCR |
| | Antisense | CCATAGTGAGCTTCTTCCTGGG | Real-time qPCR |
| ***Ibsp*** | Sense | CACTTTCCACACTCTCGGGT | Real-time qPCR |
| | Antisense | AGTTGGAGTGCCGCTAACTC | Real-time qPCR |
| ***Spp1*** | Sense | AGAAGCATCCTTGCTTGGGTT | Real-time qPCR |
| | Antisense | TCGTAGTTAGTCCTTGGCTGG | Real-time qPCR |
| ***Bglap*** | Sense | AGACAAGTCCCACACAGCAG | Real-time qPCR |
| | Antisense | GGTCAGCAGAGTGAGCAGAA | Real-time qPCR |
| ***Sox9*** | Sense | AGAGCCGGATCTGAAGAAGGA | Real-time qPCR |
| | Antisense | GCTTGACGTGTGGCTTGTTC | Real-time qPCR |
| ***Col2a1*** | Sense | TGTCCACACCAAATTCCTGTTCA | Real-time qPCR |
| | Antisense | AGGGCAACAGCAGGTTCACATAC | Real-time qPCR |
| ***Tgfβr1*** | Sense | CTCGGAACCATGAACGCTC | Real-time qPCR |
| | Antisense | GAGGATCCATCACTAGATCG | Real-time qPCR |
| ***Runx2*** | Sense | TCGGTTTCTTAGGGTCTTGGAGTG | Real-time qPCR |
| | Antisense | TGGCTTGGGTTTCAGGTTAGGG | Real-time qPCR |
| ***Tgfβ1*** | Sense | CAGCGCTCACTGCTCTTGTG | Real-time qPCR |
| | Antisense | GGGTCCCAGACAGAAGTTGG | Real-time qPCR |
| ***Col10a1*** | Sense | AGGGAGTGCAATCATGGAGC | Real-time qPCR |
| | Antisense | AGGACGAGTGGACGTACTCA | Real-time qPCR |
